# Wildlife movement and contact responses to intensive culling: implications for disease control

**DOI:** 10.1101/2025.06.13.659516

**Authors:** Kayleigh Chalkowski, Nathan P. Snow, Abigail B. Feuka, Bruce R. Leland, Kurt C. VerCauteren, Ryan S. Miller, Kim M. Pepin

## Abstract

Culling is frequently used to control animal diseases. Intensive culling can alter the movement behavior of surviving animals, especially in socially-structured wildlife species. These behavioral responses could have unexpected consequences on the spread of a disease. Thus, planning effective culling responses to diseases in wildlife hosts requires a thorough understanding of the potential impacts of culling on the target wildlife host species. We conducted a BACI design study of behavioral response to culling in wild pigs. We examined movement and contact responses in the populations using 122 GPS-collared wild pigs and three different culling methods (aerial operations, trapping, and an experimental toxic bait). Movement and contact metrics included home range area, net-squared displacement (i.e., home range shift), movement speed, distance, contact degree and contact duration. We observed increased movement distances during and after trapping treatments, and home range shifts and reduced area size after the toxicant treatment. We also observed increases in contact duration and number of unique contacts during trapping removals. Movement and contact responses varied by sex.

Our results suggest that continued, intensive culling as with extensive trapping can substantially alter wild pig space use and contact. These behavioral responses could have important consequences for disease spread when managing an introduction of transboundary animal diseases or endemic diseases.

## Introduction

Intensive culling of wildlife to achieve population reduction is frequently used to control disease transmission and reduce spread (Wobeser 2007). Some examples where culling has been attempted for disease control include rabies in wild carnivores (Rosatte 2011), devil facial tumor disease in Tasmanian devils (Lachish et al. 2010), *Mycobacterium bovis* (bTB) and Aujeszky’s disease virus in wild boar (Boadella et al. 2012), bTB in badgers (Vial and Donnelly 2012) and African buffalo (le Roex et al. 2016), chronic wasting disease in white-tailed deer (Manjerovic et al. 2014), and African swine fever in wild boar (Jo and Gortázar 2021). However, the effectiveness of culling for disease control has varied because success can depend on system-specific or local ecological factors that are not accounted for (Sokolow et al. 2019).

Unintended effects can occur when culling causes behavioral changes in the target species that alter disease spread. In socially-structured wildlife species, culling can alter conspecific interactions (Downing et al. 2023), potentially leading to changes in space use (Bielby et al. 2014, Mysterud et al. 2020), recruitment and immigration (Comte et al. 2017) or contact structure (Tosa et al. 2017) that may increase disease transmission. Evolutionary theory predicts that culling can select for lower virulence in pathogens leading to increased disease prevalence rather than control (Bolzoni and De Leo 2013, O’Neill et al. 2023). Culling can also be ineffective when disease transmission is frequency-dependent and other ecological factors exist such as population regulation through density-dependent mortality (Potapov et al. 2012) or high force of infection and continual immigration of diseased individuals (Lachish et al. 2010, Comte et al. 2017). Some culling techniques may target specific ecological groups (e.g., certain age classes or sexes), which can render culling ineffective when removing only groups that have minor roles in transmission (Streicker et al. 2012, Rogers et al. 2022). Human disturbance via intensive culling can also reduce the effectiveness of the culling effort (Ikeda et al. 2019, Keuling and Massei 2021). Taken together, ecological considerations impacting the effectiveness of culling on disease spread outcomes emphasizes the need to better understand how intensive culling impacts a targeted species before embarking on population reduction for disease control.

Population reduction can be achieved using different techniques, which may differentially affect a targeted species (Woodroffe et al. 2008, Campbell et al. 2012, Pepin et al. 2019, Snow et al. 2024c)(Campbell et al. 2010, Comte et al. 2017, Pepin et al. 2019)(Brockie et al. 1997, Sweetapple and Nugent 2009, Snow et al. 2024c) because they are. Removal techniques can also differ in the amount of effort and human presence, which could influence, potentially affecting disease transmission patterns. Thus, it is important to understand the effects of different culling techniques on population responses for planning effective disease control strategies. Wild pigs (*Sus scrofa*; typically hybrids of wild boar and domestic swine their non-native range) are a globally-distributed and abundant free-ranging species that are invasive and destructive in many countries (VerCauteren et al. 2024) and transmit a variety of diseases (Ruiz-Fons et al. 2008, Meng et al. 2009, Miller et al. 2017). Wild pigs are socially structured (Dardaillon 1988, Maselli et al. 2014, Podgórski et al. 2014, Titus et al. 2022), which can impact movement behavior (Davidson et al. 2022, Clontz et al. 2023), contact rates (Pepin et al. 2016, Podgórski et al. 2018, Yang et al. 2021), disease transmission dynamics (Pepin et al. 2021, Podgórski et al. 2022), and effectiveness of population reduction for controlling disease (Pepin and VerCauteren 2016, Fernandez-de-Simon et al. 2023). Behavioral changes of wild pigs to culling techniques are also important to understand because wild pigs are competent hosts to economically important livestock diseases such as African swine fever, Classical swine fever, Foot-and-Mouth disease, brucellosis, and bovine tuberculosis (Miller et al. 2017). For this reason, many countries have ongoing population management or outbreak response plans for management of animal disease in wild pigs (More et al. 2018, Sauter-Louis et al. 2021, Pepin et al. 2022, Brown et al. 2024). However, diseases such as bTB and African swine fever have been difficult to manage in wild pigs through population reduction once they become endemic, and require careful consideration of local disease ecology to increase effectiveness (More et al. 2018).

Much of what is known about how disease transmission in populations might respond to population reduction strategies comes from modeling because more strategies can be evaluated at lower cost (e.g., Bolzoni et al. 2014, McCallum 2016, Manlove et al. 2019, Prentice et al. 2019). Field studies that measure host population responses are important for identifying unknown processes or providing effect sizes for known processes that should be investigated by disease transmission models for identifying effective population reduction strategies (e.g., Lachish et al. 2010, Tosa et al. 2017, Mysterud et al. 2020, Downing et al. 2023). Equipping wildlife with GPS tracking devices prior to population reduction activities is a useful approach for field-based measurement of the potential behavioral responses that could impact disease transmission (Campbell et al. 2012, Tosa et al. 2017, Snow and VerCauteren 2019, Mysterud et al. 2020). We used this technique to investigate how different population reduction strategies applied to wild pigs impact movement and contact behavior. We utilized a BACI (before-after, control-impact) to provide a robust assessment of wild pig responses to population reduction by different techniques including aerial operations (i.e., shooting from aircraft), trapping, and toxic baiting.

Our experiment demonstrates the importance of evaluating multiple techniques and control intensities, and including control animals that are not exposed to the treatment for comparison. We discuss our findings in the context of planning effective population reduction campaigns for rapid elimination of a disease introduction in wild pigs – a widespread wildlife host species that are a reservoir to a wide range of diseases that impact human, domestic animal, and ecosystem health (Miller et al. 2017).

## Methods

### Study area

Our study area (225.1 km^2^) was located in the southwestern tablelands of the south central semi-arid prairies ecoregion (Bailey 1980) of northcentral, Texas, USA during January–May 2023.

The area was an active cattle ranch with a landscape dominated by shortgrass and midgrass prairies, mesquite (*Prosopis* spp.) savanna, cedar (*Juniperus* spp.), riparian areas of plains cottonwood (*Populus* spp.), and interspersed wheat fields. Temperatures averaged 7–21 °C and 97.2 mm of precipitation occurred (NOAA 2022) during our study. The topography consists of broad, rolling plains with interspersed elevated tablelands, red-hued canyons, badlands, and dissecting river breaks. Management and population control of wild pigs prior to our study was minimal. The population density of wild pigs in this region has been estimated at 2–13 wild pigs/km^2^, and reduced by an average of 55% (range = 23–90%) immediately following culling.

### Experimental design

We divided the study area into four treatment zones and randomly assigned each zone to one of three removal treatments (i.e., aerial, trapping, and toxicant), and a control (i.e., no treatment; Figure 1). Each zone ranged in area from 46.1–64.2 km^2^. We deployed GPS collars on wild pigs during Nov 2022–Jan 2023 (see detailed methodology below) throughout the 4 zones, and monitored the collared animals for 1–3 months prior to initiating treatments (i.e., pre-treatment period). We intentionally did not remove the collared wild pigs during the population control treatments. We left these animals on the landscape to represent survivors of our activities and the simulated TAD. The only exception was for the toxic baiting treatment where we could not control which wild pig consumed the toxic bait.

**Figure 1.**
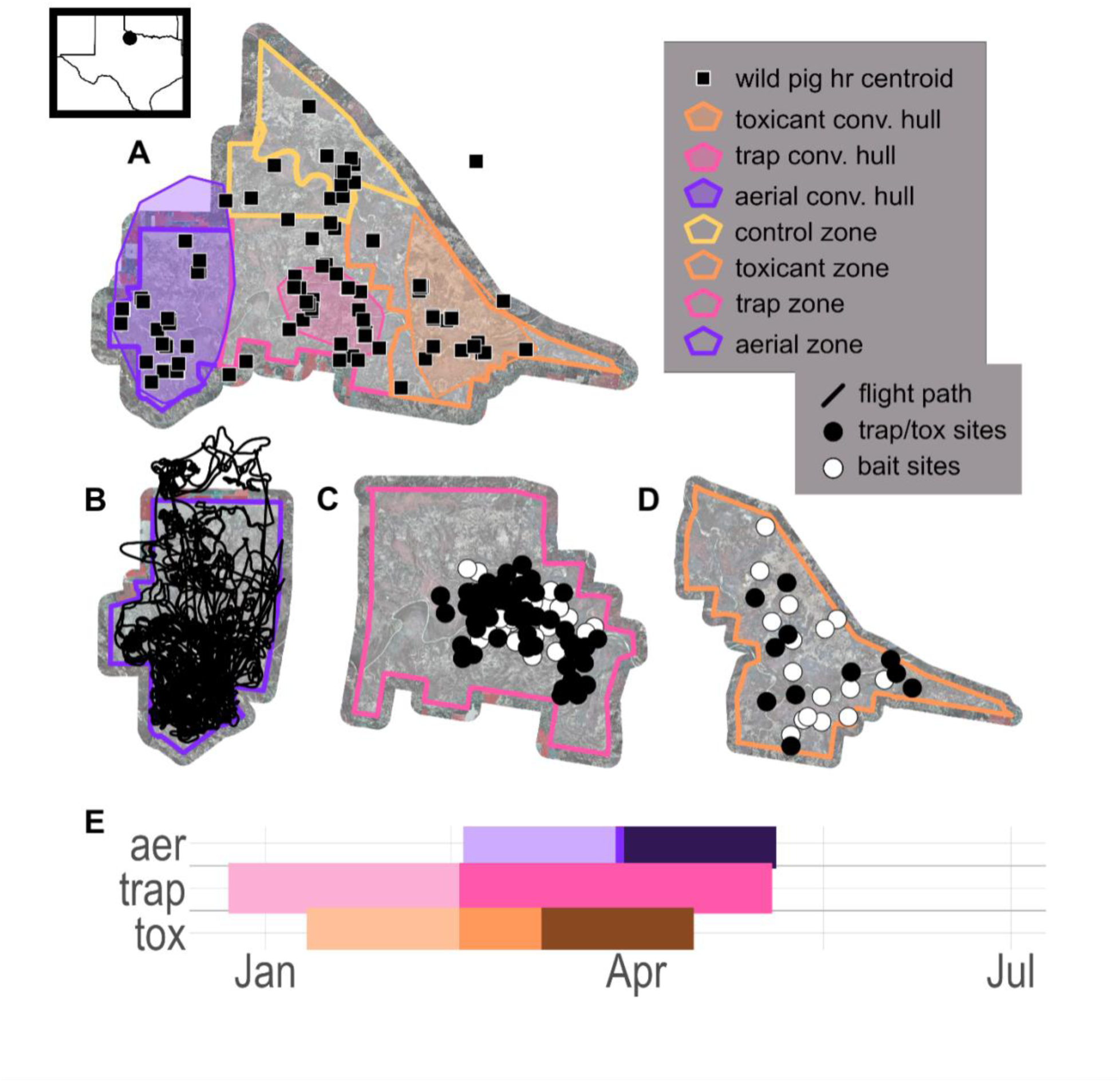
A) study area with removal treatment and control zones (thick border polygons with no fill); convex hulls for each treatment (thin border polygons with color fill); and site locations for wild pig collaring captures (white squares). B) aerial zone, with flight paths, trimmed to include only paths with intensive aerial gunning operations; C) trapping zone, pre-baiting sites (white circles) and trap deployment sites (black circles); D) toxicant zone, with pre-baiting sites to assess wild pig visitation (white circles) and toxicant deployment sites (black circles); E) timeline of removal activities for each strategy; aer-aerial control operations; trap-trapping removals; tox-toxicant removals.

### Deploying GPS collars

We attached GPS collars (Vertex Lite, VECTRONIC Aerospace GmbH, Berlin, Germany) on adult wild pigs >35kg. Specifically, we deployed collars on 1–3 animals per adjacent social group in attempt to distribute the collars across a contiguous and adjacent network of wild pigs within each treatment zone. We programmed the collars to collect geolocations every 20 minutes (15 minutes in the toxic baiting zone). Overall, we deployed 122 collars across the study area.

All collars were set to drop off on 01 July 2023 (except 01 May 2023 in the toxic treatment zone). All research methods were approved by the USDA National Wildlife Research Center, Institutional Animal Care and Use Committee (QA-3311 and QA-3470).

### Population Control Treatments

Detailed methodologies for conducting culling of wild pigs for each treatment were described in Snow et al. (2024b), Chalkowski et al. (2024), and Snow et al. (2024a). In brief, for the trapping and aerial operations we generated focal concentration areas by viewing satellite geolocations from the collars and centering our removal activities in areas where collared pigs were located. For toxic baiting and to meet the purposes of also evaluating and experimental toxic bait, we focused on removing wild pigs within the overall treatment zone and did not generate a focal area.

Overall, we culled 296 wild pigs with trapping, 256 with aerial operations, and an estimated 58 with toxic baiting. For toxic baiting, we estimated the number of wild pigs culled by walking transects and locating carcasses, and examining remote camera imagery to identify wild pigs that consumed toxic bat and never returned to the bait sties (Snow et al. 2024b). Immediately following the treatments, Snow et al. (2024b) reported declines in the wild pigs population of 89.9% in the trapping zone, 51.8% in the aerial operations zone, and 23.3% in the toxic baiting zone.

### Treatment assignments

All data formatting and analyses were conducted in R version 4.3.2 (R Core Team 2023).

We assigned all animals to 1 of the 3 treatments, or control, based on their space-use. We determined the spatial area of the removal activities by determining the convex hull around all the removal activities for each treatment, respectively (i.e., bait and trap sites in the trapping zone, helicopter track logs in the aerial operation zone, and bait and toxic sites in the toxic baiting zone), using the sf R package (Pebesma 2018). Then, we generated minimum convex polygon (MCP) core areas (50% MCP) and home ranges (95% MCP) for each of the collared wild pigs using the amt R package (Signer et al. 2019). We assigned a wild pig to the one of the 3 treatments if its 50% core area overlapped with any of the respective treatment boundaries (Figure 1, Supplementary Figure 1). We designated control animals as any wild pig whose 95% home range did not overlap any of the treatment boundaries (Supplementary Figure 1).

### Geolocation data processing

We processed geolocations by trimming incomplete days and any obviously erroneous points (e.g., maximum speed of 25 km/h for any consecutive points >50 m apart; Mayer 2009 and Gupte et al. 2021). We then filtered geolocation data for the overall dataset to ensure matching durations of each removal period (Figure 1), and according to specifications for each movement response (Supplementary Figure 1) including range residency requirements for home range area calculations (Silva et al. 2022) and fix rates >20 minutes for movement responses requiring continuous time movement models (CTMM)(Yang et al. 2023). We assigned geolocations to treatment period ‘before’, ‘during’, or ‘after’ according to respective removal activity dates (Figure 1, 3, 4).

We tested sensitivity of movement responses home range area and NSD (Table 1) to lower fix rates. We found that all area estimates for both 20 and 240 minute subsampled datasets were within the 95% confidence intervals of the area estimated from the full dataset (Supplementary Figures 6, 7). For displacement, the maximum difference in displacement was not biologically significant (Supplementary Figures 8, 9). Thus, we kept data from individuals with lower fix rates (Supplementary Figure 2). Several individuals in the toxicant treatment did not survive toxicant bait treatment. Thus, we tested whether the sensitivity of our model parameters to removal or inclusion of these individuals. We found that model parameter estimates were not biased by inclusion of individuals that succumbed to toxicant bait (Supplementary Figure 12).

**Table 1.**
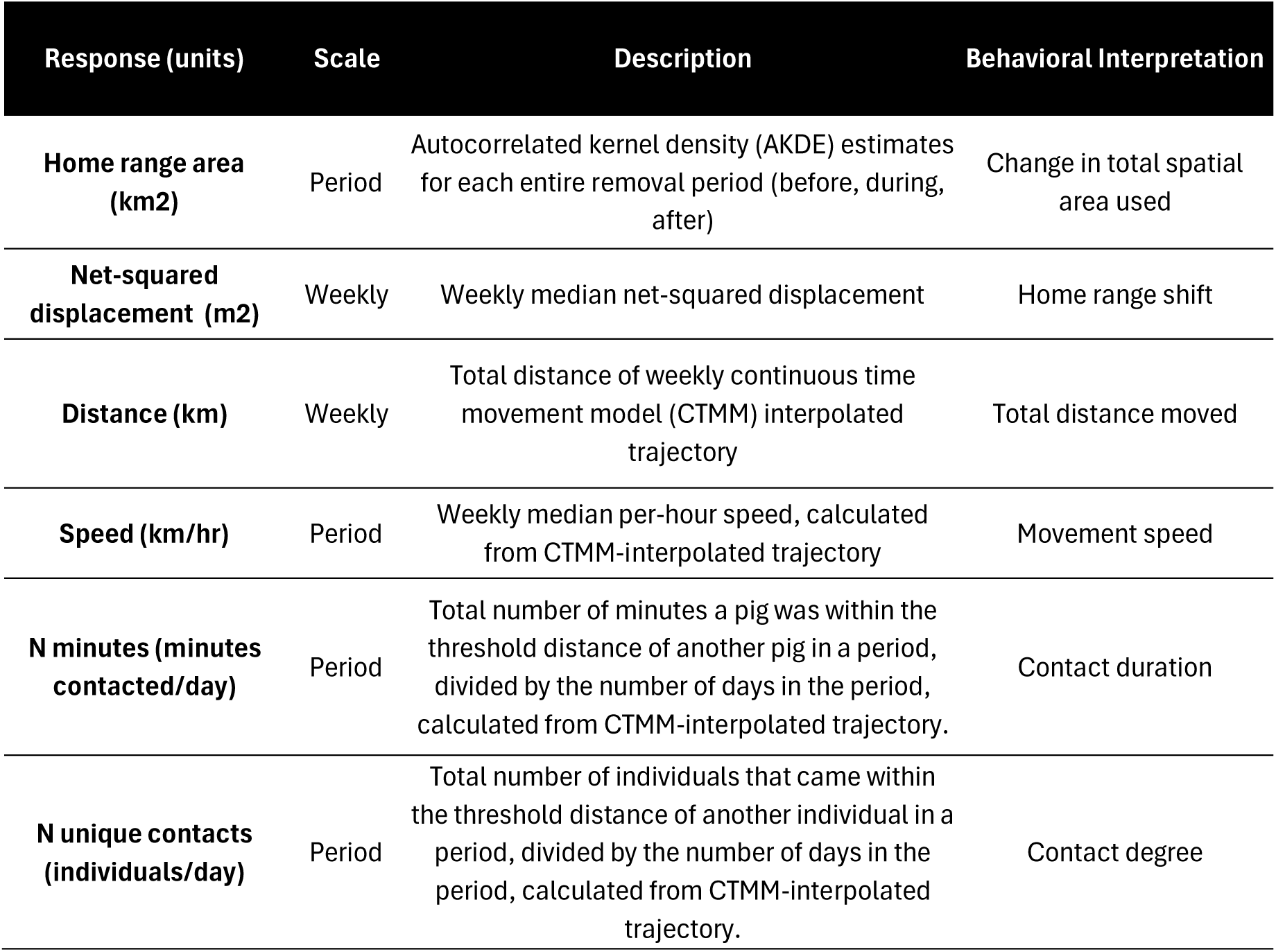
Movement and contact responses analyzed in response to removal operations (aerial control, trapping, toxicant) in wild pigs.

| Response (units) | Scale | Description | Behavioral Interpretation |
| --- | --- | --- | --- |
| **Home range area (km<sup>2</sup>)** | Period | Autocorrelated kernel density (AKDE) estimates for each entire removal period (before, during, after) | Change in total spatial area used |
| **Net-squared displacement (m<sup>2</sup>)** | Weekly | Weekly median net-squared displacement | Home range shift |
| **Distance (km)** | Weekly | Total distance of weekly continuous time movement model (CTMM) interpolated trajectory | Total distance moved |
| **Speed (km/hr)** | Period | Weekly median per-hour speed, calculated from CTMM-interpolated trajectory | Movement speed |
| **N minutes (minutes contacted/day)** | Period | Total number of minutes a pig was within the threshold distance of another pig in a period, divided by the number of days in the period, calculated from CTMM-interpolated trajectory. | Contact duration |
| **N unique contacts (individuals/day)** | Period | Total number of individuals that came within the threshold distance of another individual in a period, divided by the number of days in the period, calculated from CTMM-interpolated trajectory. | Contact degree |

**Table 2.** Removal methods (aerial, trapping, toxicant) with movement and contact responses, in the context of disease management implications, and further disease management research needs.

| Removal Method | Response | Disease management implications | Research Needs |
| --- | --- | --- | --- |
| Aerial | decreased speed and distance moved for females, no changes in contact | For short durations of aerial control (~3 days), few impacts on animal movement and contact. | Need to understand movement and contact responses to longer durations of intensive aerial population control (>3 days) |
| Trapping | increased distance moved and reduced home range shifts for females; increased distance and increased home range shifts for males; increased contact duration and number of unique contacts during trapping for both sexes; reduced contact duration and degree after trapping | Females are less likely to move away from the removal zone and spread the disease across the landscape, but females live in larger groups that contact more individuals. Adult males may contact fewer individuals, but can shift home range in response to removal activities and pose challenging for eliminating the disease on the landscape. Managers should still prioritize whole-sounder removal to reduce pigs on the landscape overall, but take advantage of opportunities to capture lone wild boars. | Whole sounder removal techniques well-studied (Lavelle et al. 2021), but methods for targeting lone boars are needed. |
| Toxicant | increased distance moved and reduced home range shifts and home range area for females; spurious home range shifts in males |  |  |

### Movement response calculations

We calculated four movement and two contact responses (Table 1, Figures 2, 3). For home range area, we used continuous time movement (CTMM) models and autocorrelated kernel density estimation (AKDE)(Calabrese et al. 2016) using the ctmm R package (Calabrese et al. 2016). For AKDE, we evaluated for any home range estimates for wild pigs during any period that resulted in a >3% order of bias (Silva et al. 2021) using R package segclust2D (Patin et al. 2019). For these, we separated tracks into clusters comprised of divergent behavioral states (Silva et al. 2021). We excluded any tracks that could not be segmented into clusters with <3% (i.e., did not meet range residency requirements, Silva et al. 2022, Supplementary Figure 1). We calculated median net-squared displacement (NSD) by calculating the distance between weekly median geolocations. Speed and distance were calculated using the continuous-time movement models (CTMMs) interpolated to one-minute intervals for all wild pigs along their trajectories. Using these predicted trajectories, we calculated weekly distance moved by summing the total weekly distance and speed of the trajectory for each animal, each week/treatment period.

**Figure 2.**
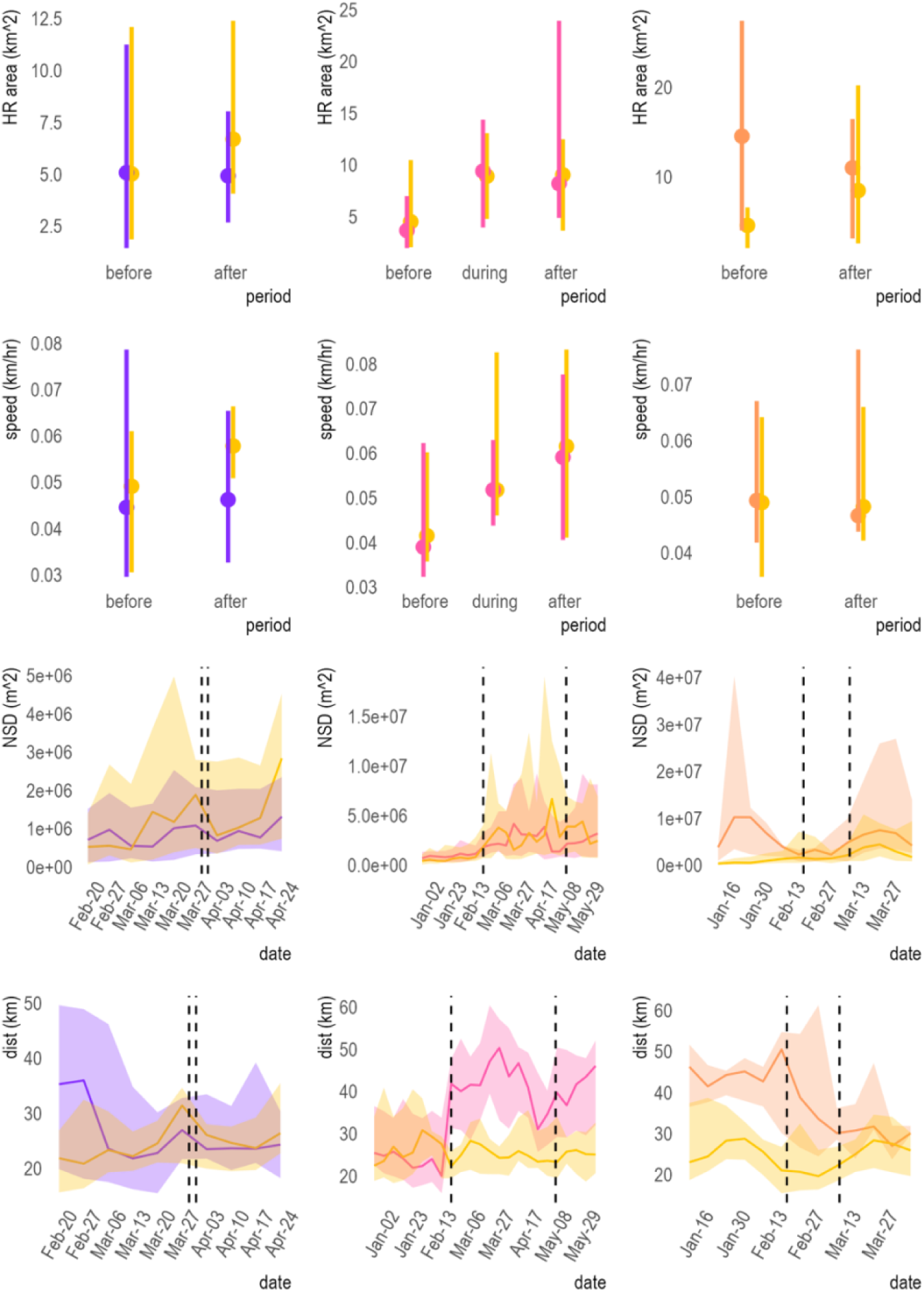
Filtered and summarized raw movement response data summarized by each removal period including before removal activities occurred, during each removal treatment; and after removal activities ended. Movement responses summarized with dot-whiskers (HR area and speed) were summarized across each entire removal period, with respect to the removal method. Movement responses summarized with line graphs (NSD, distance) were summarized on a weekly scale. Dashed lines on line graphs indicate the start and end of removal activities, respective to each removal method. HR area-total home range area for each period; speed-median speed for each removal period; NSD-weekly median net squared displacement; dist-total distance of movement path per week. yellow-control; purple-aerial removals; pink-trapping removals; orange-toxicant removals

**Figure 3.**
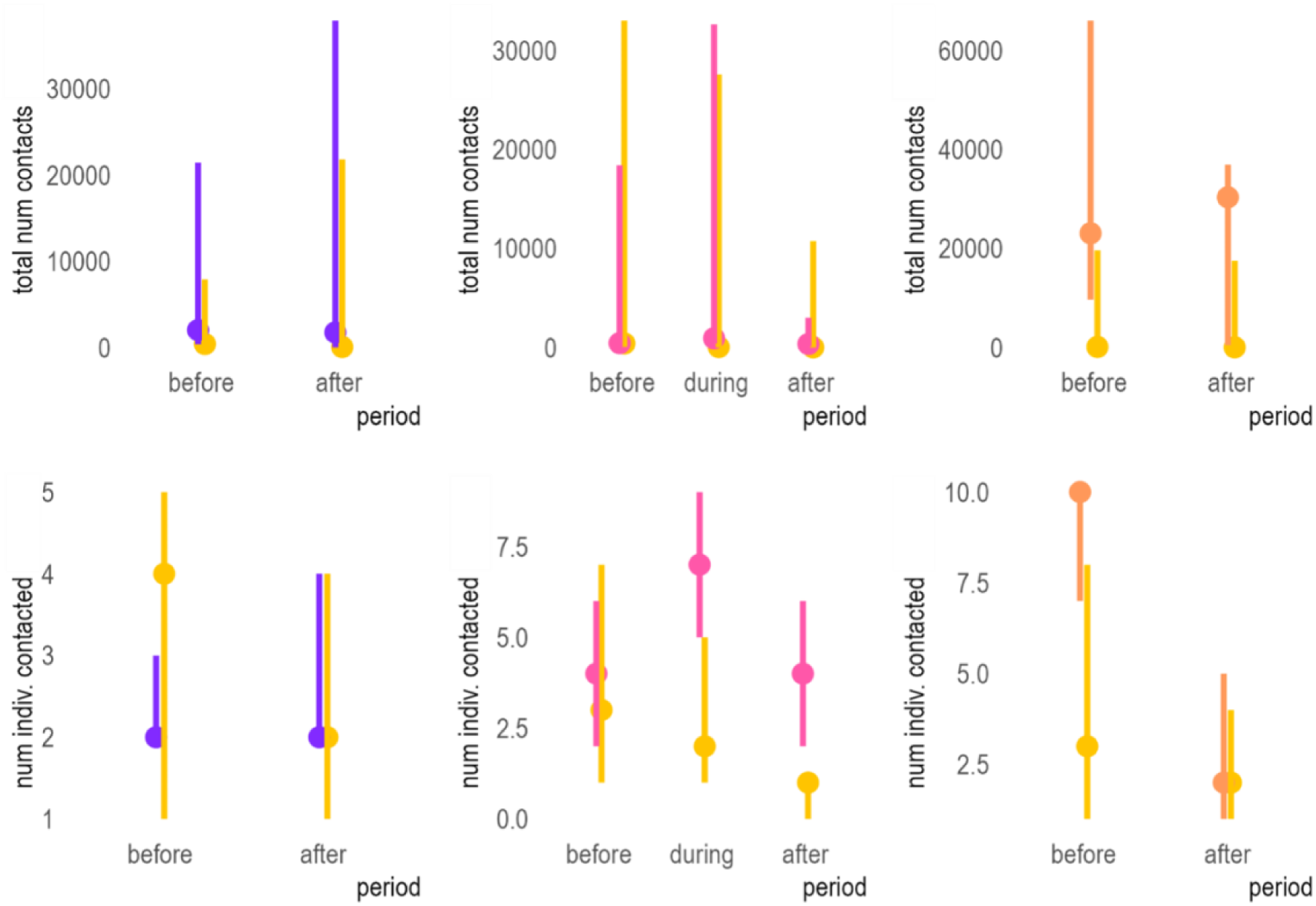
Filtered and summarized raw contact response data (total number of contacts per week; number of individuals contacted per day) by each removal period including before removal activities occurred, during each removal treatment; and after removal activities ended. All contact data was summarized across each entire removal period, with respect to the removal method. yellow-control; purple-aerial removals; pink-trapping removals; orange-toxicant removals

To calculate our contact metrics (contact duration in minutes/day and number of unique individuals contacted (Table 1)), we simulated one-minute trajectories that matched the timestamp of each pig for all other pigs in the treatment area (Yang et al. 2023). For each minute timestamp, pairwise distances <10 m were classified as contacts. The contact duration was the sum of instances the focal individual was within threshold distance of another pig, and the contact degree was the number of unique pigs the focal individual came into contact with. To mitigate computational overhead, we only analyzed pairwise distance between pigs in the same treatment group whose 95% MCPs overlapped by 1% or more in area. We calculated MCPs using the amt R package (Signer et al. 2019). We only analyzed survivors of removal treatments (as reference pigs or contacts), as including them would negatively bias estimates of contact between removal periods (Supplementary Figure 1).

### Statistical Analyses

We examined the influences of each treatment type on movement and contact for wild pigs using a before-after-control-influence (BACI) design. We compared animals from each treatment type to the control animals separately, because all treatment periods did not overlap across treatments (Figure 1). We used generalized mixed linear models with a gamma distribution, using the glmmTMB package (Brooks et al. 2017) in R to evaluate for changes in the movement and contact responses among periods. To test the effect of each removal treatment on movement and contact, we ran two sets of models for each response and removal type, one with the fixed parameters of period + treatment + period × treatment, and the other period + treatment + period × treatment × sex. (add equations below) We included wild pig ID as random effects to account for repeated measures and individual-level variation.

We tested for temporal autocorrelation for weekly parameters (NSD, movement distance, and speed) using Durbin-Watson (DW) test and considered residuals significant temporal and spatial autocorrelation where p<0.05. Models with evidence of significant temporal autocorrelation were refit with an autoregressive error structure, AR(1), of week nested within each pig identification number to account for temporal autocorrelation. We also tested each model for spatial autocorrelation with Moran tests, using mean location of each pig across week or period (depending on scale at which response was evaluated). Any models with significant spatial autocorrelation were refit with a spatial autocorrelation error structure with either glmmTMB or sdmTMB (Brooks et al. 2017, Anderson et al. 2024). Lastly, for any model with interactions (treatment * period) that had large effect sizes (>20), but were no longer significant with the additional of sex as a covariate, we conducted a leave-one-out to identify whether large effect size are the result of overall population trends, or due to high inter-individual variation in responses.

## Results

We observed significant effects of removals on several movement responses (Figure 4; Supplementary Figures 2, 3, Supplementary Tables 1-4), where all results reported from these models are described relative to control individuals. We observed significant reductions in speed after aerial removals (-10m/hr difference, p=0.032, Figures 5, 6) and increases in distance moved during (+16.7 km difference, p=6.3e-21, Figure 7, Figure 5) and after (+9.3 km difference, p=8.7e-8) trapping. We saw increased distance moved during toxicant deployment (+7.46 km difference, p=4.0e-4, Figure 5, 6), and increased home range shifts (+4.81 km difference, p=4.1e-6) and reduced home range area changes (-10.4 km difference, p=0.0076) after toxicant deployment.

**Figure 4.**
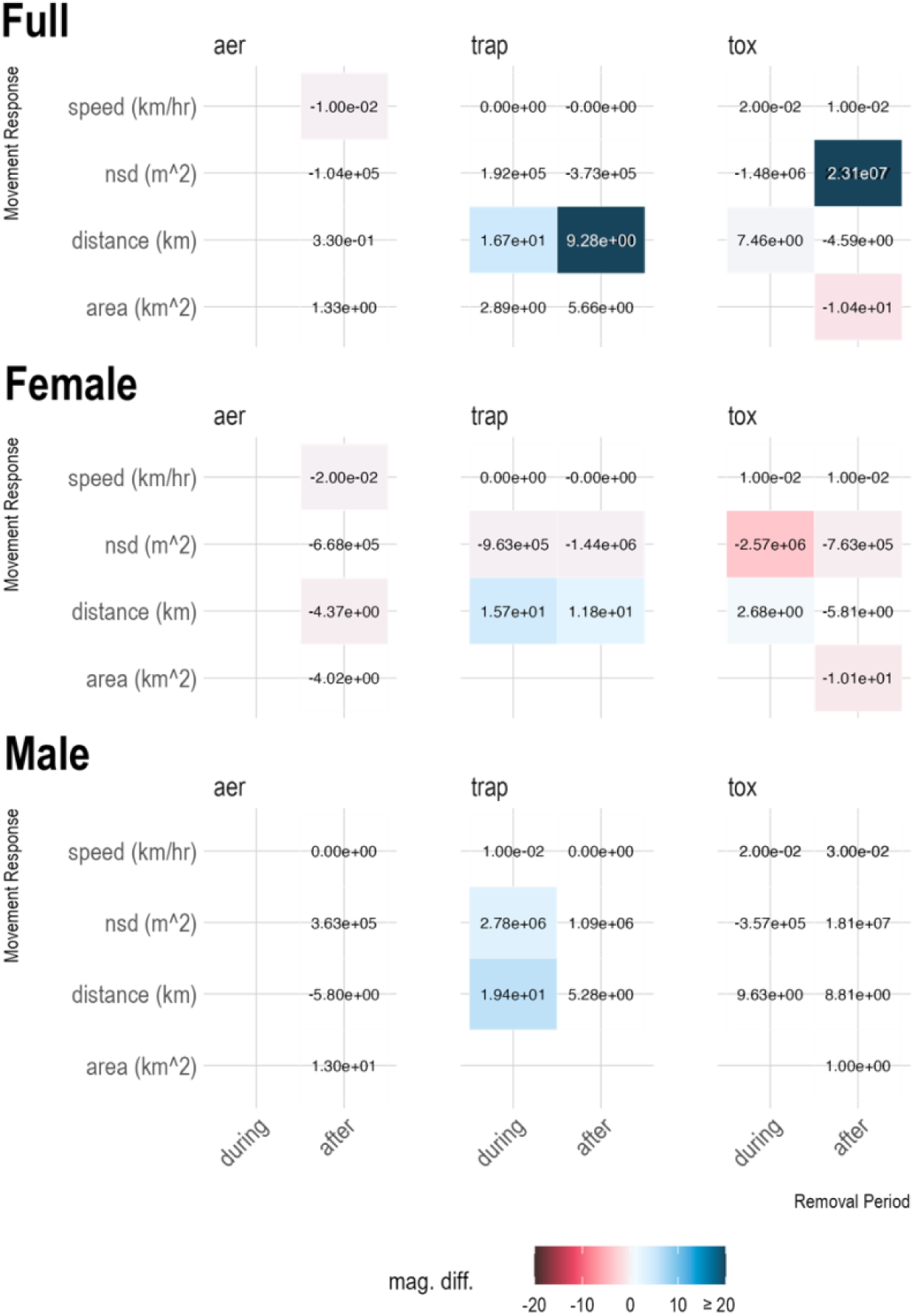
Heat map with text labels indicated predicted changes for each movement response, relative to controls. These values were calculated by subtracting predicted changes in movement, for each response, of treatment samples across removal periods from predicted changes in movement of control individuals (i.e., the change in movement of treatment). The color scaling reflects the absolute magnitude of difference between treatment and control movement changes, where negative values (red) indicates a reduction for each movement response. Aer-aerial operations removals; tox-toxicant removals; trap-trapping removals. area-total home range area for each period; speed-median speed for each removal period; nsd-weekly median net squared displacement; distance-total distance of movement path per week.

**Figure 5.**
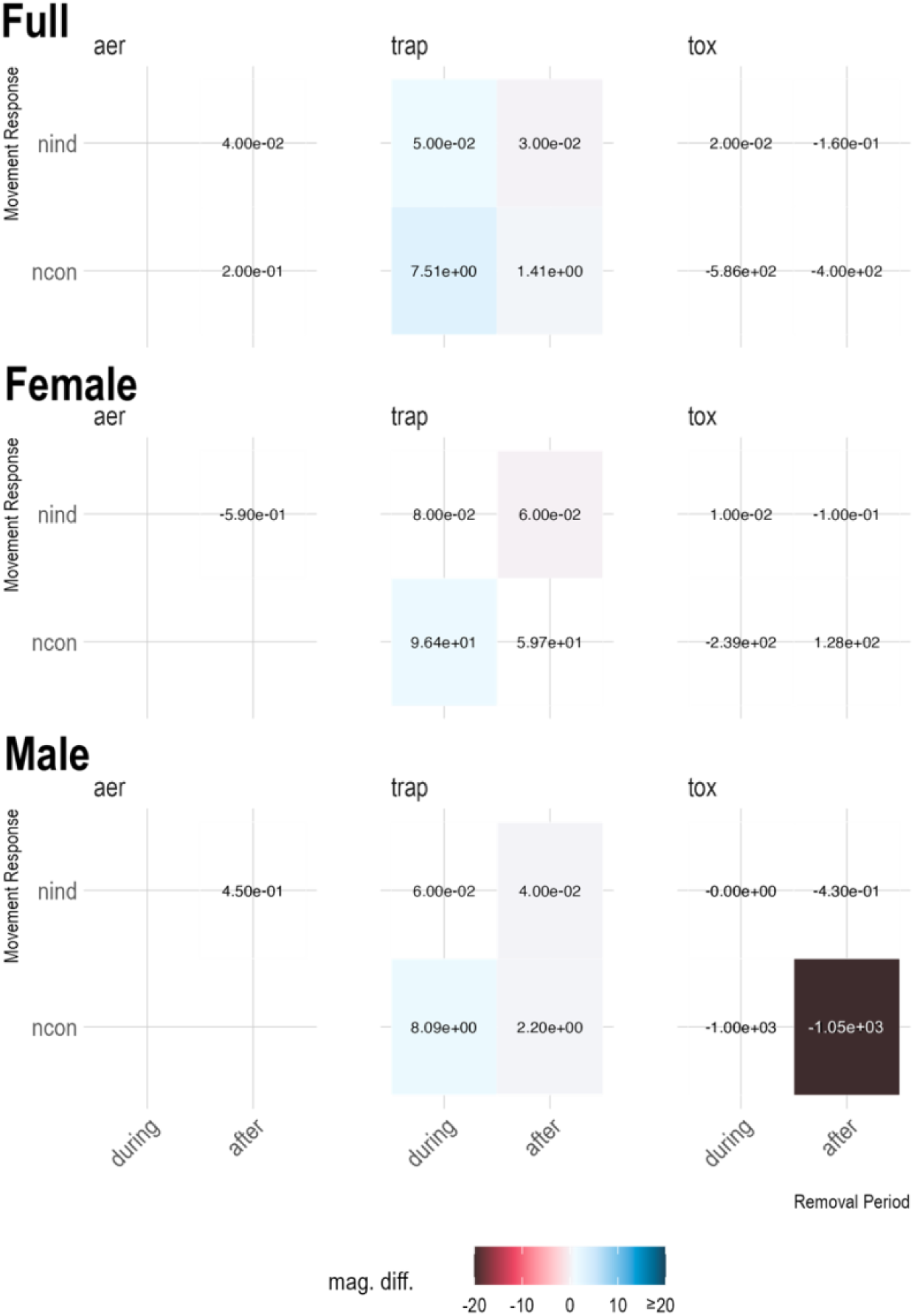
Heat map with text labels indicated predicted changes for each contact response, relative to controls. These values were calculated by subtracting predicted changes in movement, for each response, of treatment samples across removal periods from predicted changes in movement of control individuals (i.e., the change in movement of treatment). The color scaling reflects the absolute magnitude of difference between treatment and control movement changes, where negative values (red) indicates a reduction for each movement response. Aer-aerial operations removals; tox-toxicant removals; trap-trapping removals. Ncon-total number of contacts per day; Nind-number of unique individuals contacted per day.

Trapping was the only removal treatment that had significant effects on contact among pigs when sex was not considered (Figure 5, Supplementary Figure 4). Contact duration increased during (+39.0 minutes/day difference, p=9.0e-3) and after (+16.1 minutes/day difference, p=0.02, Figure 8, 10) trapping occurred. Contact degree also increased during (+0.05 individuals/day difference, p=0.02, Figure 8, 10) and after (+0.03 individuals/day difference, p=2e-3, Figure 8, 10) trapping.

We observed differences in movement responses by sex (Figure 4, Supplementary Figure 3). Female reduced movement rates: reduced speeds and distances moved following aerial removals, reduced home range shifts, reduced home range shifts during and after both trapping and toxicant deployments, and reduced home range area after toxicant deployments. We did not observe any reductions in movement for males. We observed increased home range shifts of males (i.e., nsd) relative to females during trapping deployment (p=3.9e-4), such that males shifted more (+1.7km difference, p<0.001), and females shifted less (-0.98 km difference; p=0.001) during trapping deployment.

Contact only differed significantly by sex for toxicant and trapping treatments (Figure 5, Supplementary Figure 5). Male pigs had substantially lower contact durations per day after toxicant treatment (-1190 minutes/day difference, p=0.01). Male pigs made more contacts per day (+2.20 minutes/day difference, p=0.01) after trapping. Female pigs made substantially more contacts per day (+96.4 minutes/day difference, p=0.04) during trapping, as well as slightly more unique individuals contacted per day after trapping (+0.06 individuals/day difference, p=0.03).

No significant sex effects were observed for the number of total contacts made per day by pigs during or after aerial operations. In our leave one out analysis for two models with high (>20) parameter estimates, we found that the interaction between toxicant removal treatment and removal period for home range area was sensitive to removal of two individuals (Supplementary Figure 10). The leave-one-out analysis for the interaction of trapping treatment and removal period for change in distance was robust to removal of any individual (Supplementary Figure 11).

## Discussion

### Movement and contact responses to removals vary by removal method

It is notable that present and previous evaluations of aerial removal treatments (Saunders and Bryant 1988, Dexter 1996) have observed little to no movement changes under aerial removal operations despite considerable population reductions. For example, in the present study, we reduced the population by 52% (5 pigs/km2) by aerial removals. Earlier studies conducted greater numbers of removals and/or over slightly longer durations (5 days/7.9 pigs/km2 (Saunders and Bryant 1988); 0.3 pigs/km2, 4.5 days (Dexter 1996)), and still observed negligible differences in movement of wild pigs. In fact, the number of individuals removed under aerial treatments in our study was similar to the number of removals in our trapping treatments (Figure 1). It is also notable that toxicant removals removed a much smaller number of individuals, but still had significant effects on movement and contact. There could be a few non-mutually exclusive interpretations of this pattern. For one, perhaps longer durations of disturbance are required for behavioral changes in wild pigs. Pigs are well known to learn cooperatively from conspecifics, and have spatial memory (Morelle et al. 2024), so perhaps experiences with removal operations take effect at larger scales as learned behaviors over time. Alternatively, it could be that the magnitude of population reduction is less important than the disturbance imposed on the population by removal operations. For example, wild boar responses to types of disturbances vary (Faltusova et al. 2024), suggesting that the observed movement response differences in our study could be due to pigs perceptions to the different operations.

Although aerial operations still involved human presence of ground crews to locate carcasses, (Snow et al. 2024) these did not involve the same operational activities as trapping and toxicant deployments. In our trapping design, we implemented a dense trapping grid (Snow et al. 2024) that was regularly maintained with cameras to assess pig visitation, building of traps and other equipment when known pig presence. This dense trapping grid which involved regular maintenance of bait sites to regularly assess pig presence, in fact, could explain the increase in contact rates for this modality compared to earlier studies (Campbell et al. 2012, Bastille-Rousseau et al. 2021, Fischer et al. 2016). Toxicant deployment also may impose a higher level of disturbance, as we observed changes in movement responses similar to earlier studies of toxicant removals on social species (Sweetapple et al. 2009). It is operationally similar to trapping (Chalkowski et al. 2025) in that it requires pre-baiting to identify pig presence, setting up of bait stations which require acclimation, and operation over longer durations compared to aerial control. Additionally, deployments of our experimental toxicant bait also required transect surveys by personnel every 50 m on 400x400m grids to identify wild pigs that succumbed to toxicant bait, which could have represented strong disturbance on wild pigs.

Further work to evaluate the effects of different removal methods on wild pigs should compare movement response changes at finer scales in relation to specific operational activities within a BACI design. For example, observing specific removal operations (i.e., baiting a trap, conducting a transect survey) on wild pig movement responses within the same time period.

Additionally, understanding the threshold of duration at which point movement responses start occurring may provide guidance for operations to spatially optimize activities to avoid inducing counterproductive behavioral changes.

### Movement and contact responses to removals vary by sex

We observed similar responses between sexes for contact, but differences in movement responses. Females frequently reduced movements in response to removal activities (Figure 7), with the exception of increased movement distances during trapping removals observed for both males and females. This similarity of increased distances under trapping and toxicant is similar to previous findings on wild pig responses to baiting, such that males and females responded similarly to baiting with increased movements (Snow and Vercauteren 2019). Reductions in movement responses for females could be explained by increased vigilance behavior in females (Davidson et al. 2021). It is also notable that the two individuals that strongly influenced some of our findings for toxicant deployment movements were both males (Figure S5). Males move more than females (Friesenhahn et al. 2023, Cavazza et al. 2024), and increases in home range may be driven by access to females, whereas females tend to follow resources (Cavazza et al. 2024).

Thus, perhaps the large movement changes observed for these males were in response to removals of nearby females.

We observed a drastic decrease in the contact durations by male pigs after toxicant deployment. We limited our analysis of contact to only individuals that survived the entire trial, meaning the decrease in contact is not a result in animal death. Rather, individuals that previously moved together ceased doing so after toxicant deployment, suggesting sounder structure may have changed. The reverse effect was observed after trapping, with greater contact rates overall. The increase in distance moved and decrease in NSD for female pigs suggests females moved more within a smaller area over the trapping deployment, which could contribute to increased contact. Males exhibited increased movement with increased NSD, which could keep contact rates similar throughout the study.

### Movement and contact under a disease control operation

Each of the movement and contact changes we observed may have different outcomes in the context of a disease control operation. The increased home range shifts we observed in particular may affect the spatial spread of a disease (Podgorskiand Smietanka 2018) and outbreak size (Beninca et al.2020, Keeling et al. 2001 and may require implementing control over a larger spatial area (Lintott et al. 2013, Pepin et al. 2022). Additionally, we observed that it may be a few individuals driving the home range shift effect, particularly for toxicant deployments. Movements of a population characterized by generally localized movements with occasional large dispersals can drive the spatial spread of disease (Beninca et al. 2020, Keeling et al. 2001).

We also observed reduced home range areas, which could benefit disease control operation success. There is less definitive evidence that home range area can affect spatial spread or outbreak size of a TAD (Podgorski and Smietanka 2018). However, increased home range size can make disease control operations such as vaccine deployment more difficult due to hosts being more spatially spread (McClure et al. 2020). We didn’t observe any meaningful changes in movement speed, contrary to previous studies (Mysterud et al. 2020). However, we did find that both males and females moved increased distances overall during and after trapping, and during toxicant deployments. Movement distance is distinct from home range area and net-squared displacement. Compared to home range and displacement characterizations, understanding path-level transmission rates can reveal inter-individual heterogeneity in transmission, which can be meaningful for epidemic outcomes (Wilber et al. 2022). These fine scale, individualized differences in transmission as a result of changes in the movement paths may be relevant for disease control operations.

Lastly, the increase in contact duration and number of unique individuals contacted we observed during and after trapping removals is important for disease control operations, because contact rate influences epidemiological outcomes for any system involving direct or indirect transmission (Hu et al. 2013, Craft 2015, Chen et al. 2014). Our findings were contrary to previous findings for contact duration (Tosa et al 2017). Increase in new individuals contacted is similar to previous findings for contact structure changes (Downing et al. 2023, Allen et al. 2022).

Future research to connect these movement and contact response findings to disease control are needed. For one, we know that many of these movement and contact responses evaluated are epidemiologically relevant (Podgorski and Smietanka 2018, Beninca et al.2020, Keeling et al. 2001, Pepin et al. 2022), but there is less certainty around the sensitivity of each of these movement and contact changes to epidemiological outcomes and disease control operations, and which changes are more meaningful for outbreak dynamics such as probability of outbreak die-out or spatial spread. Concurrently, studies that also evaluate these parameters in alignment with disease control operational parameters (e.g., control area size, time to detection) needed for elimination of a pathogen are needed to ensure elimination under culling operations. Understanding how the magnitude of the responses observed can translate to an outbreak scenario can allow for development of targeted strategies to mitigate behavioral responses which are most detrimental to disease control success.

## Acknowledgments

The research was supported by NIFA-AFRI Agricultural Biosecurity program (GRANT13428587) and the United States Department of Agriculture. The findings and conclusions in this publication are those of the authors and should not be construed to represent any official US Government determination or policy. Mention of commercial products or companies does not represent an endorsement by the US government. The authors declare no conflicts of interest related to this work. We thank H. Ownbey and G. Studdard for access to private property. We thank Animal Control Technologies Australia for providing the placebo and SN-toxic bait. We thank multiple people from Texas Wildlife Services and Texas Parks and Wildlife for assisting with data collection. We thank anonymous reviewers for their comments on this manuscript.

