## Supplements for "Wildlife movement and contact responses to intensive culling: implications for disease control"

**Supplementary Figure 1.** Flowchart depicting process for filtering 122 original collars and 1,474,480 geolocations for each movement response analysis and removal method. Purple- aerial removals; pink- trap removals; orange- toxicant removals; Trt- treatment; Ctrl- control; Area- total home range area for each period; speed- median speed for each removal period; NSD- weekly median net squared displacement; distance- total distance of movement path per week; ncon- total number of contacts per day; nind- number of unique individuals contacted per day.

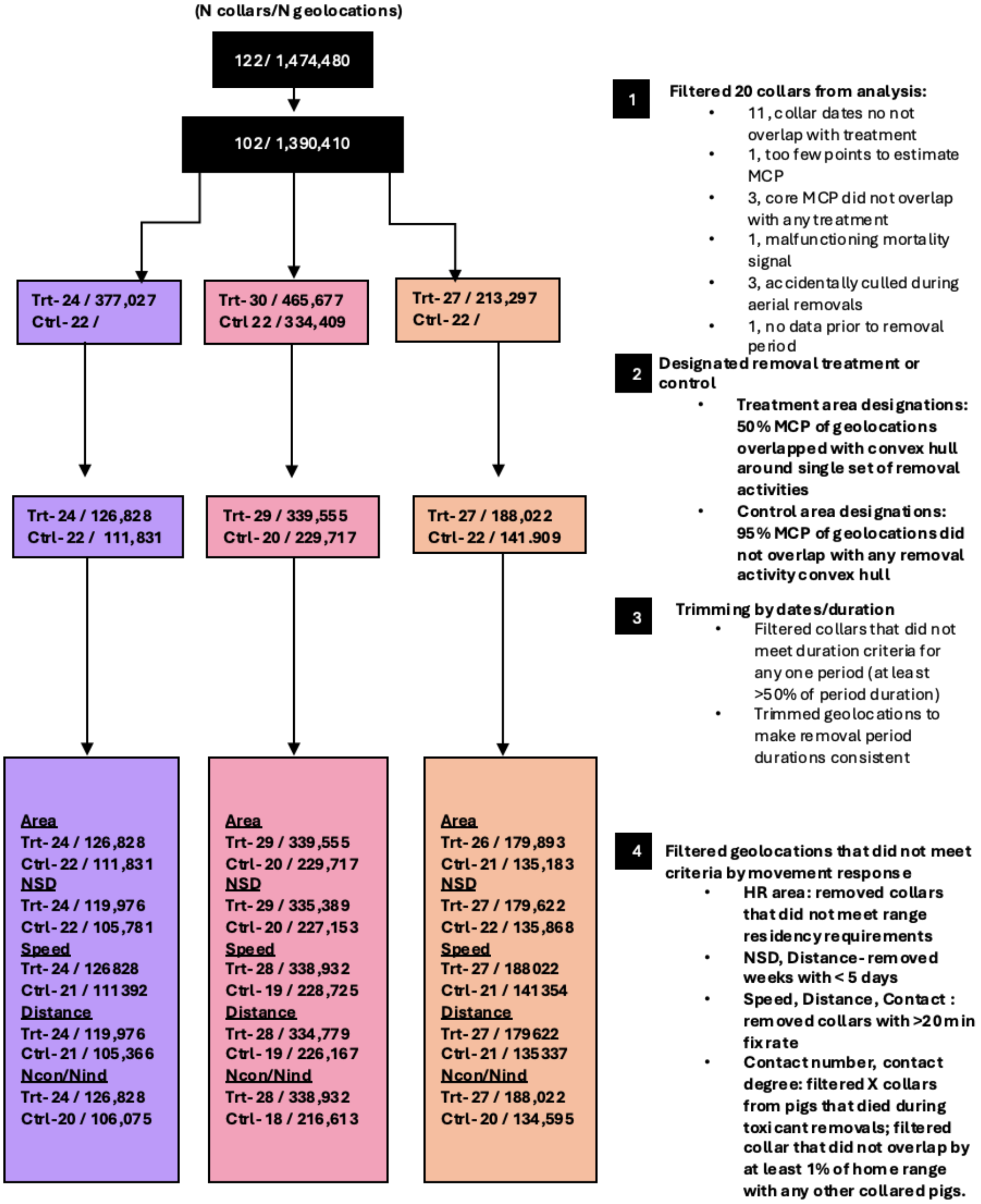

**Supplementary Figure 2.** Model predictions for movement responses by removal period (before- before removal activities occurred; during- during active removal activities; after- after removal activities ended), respective to each removal method. *- interaction between removal treatment and removal period (i.e., effect of removal relative to control) was significant. Error bars indicate 95% confidence interval. area- total home range area for each period; speed- median speed for each removal period; NSD- weekly median net squared displacement; distance- total distance of movement path per week. yellow-control; purple-aerial removals; pink-trapping removals; orange-toxicant removals

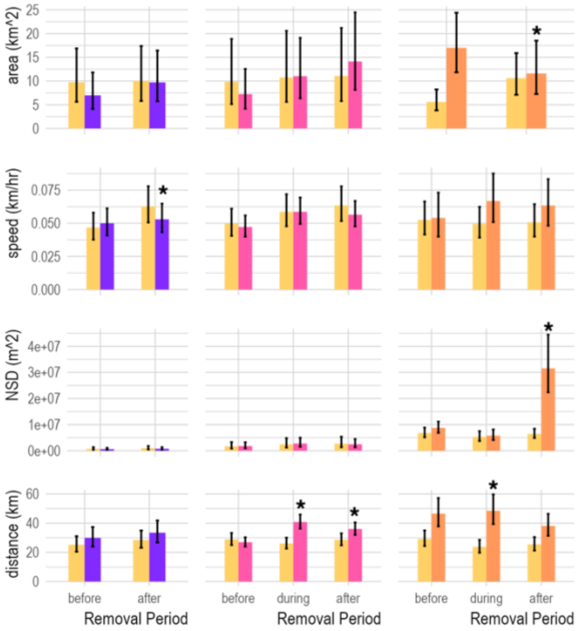

**Supplementary Figure 3.** Model predictions for movement responses by removal period (before- before removal activities occurred; during- during active removal activities; after- after removal activities ended), respective to each removal method, including the interaction of sex. *-interaction between removal treatment and removal period (i.e., effect of removal relative to control) was significant. **-interaction between removal treatment, period, and sex was significant. Error bars indicate 95% confidence interval. HR area- total home range area for each period; speed- median speed for each removal period; NSD- weekly median net squared displacement; dist- total distance of movement path per week. yellow-control; purple-aerial removals; pink-trapping removals; orange-toxicant removals

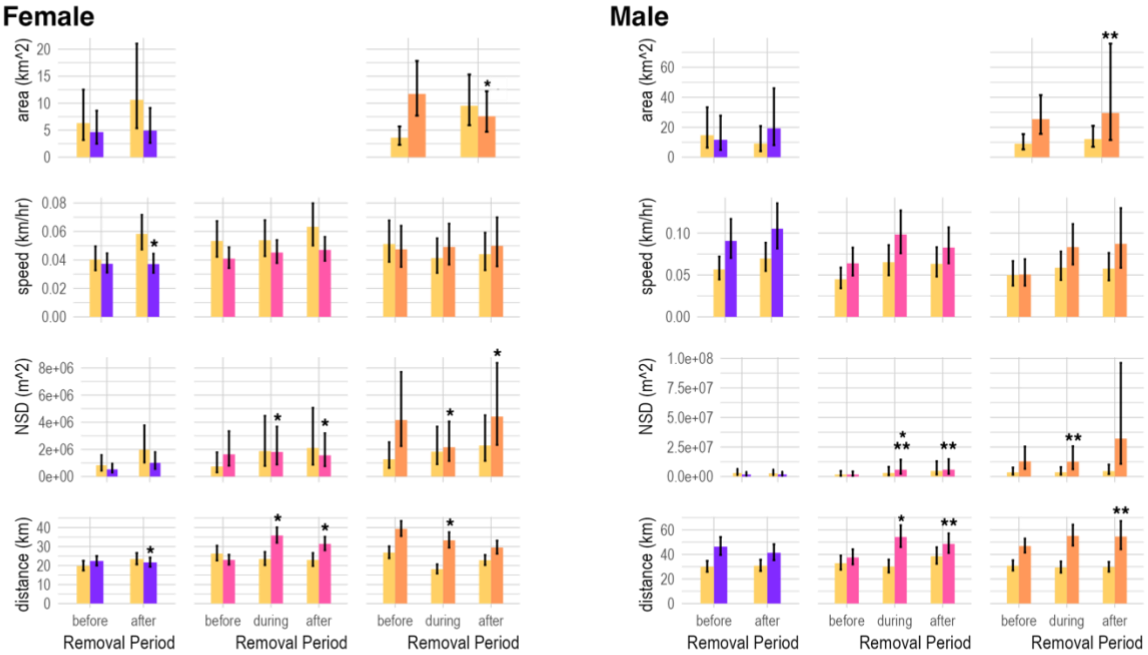

**Supplementary Figure 4.** Model predictions for contact responses by removal period (before- before removal activities occurred; during- during active removal activities; after- after removal activities ended), respective to each removal method. *- interaction between removal treatment and removal period (i.e., effect of removal relative to control) was significant. Error bars indicate 95% confidence interval. Ncon- total number of contacts per day; Nind- number of unique individuals contacted per day. yellow-control; purple-aerial removals; pink-trapping removals; orange-toxicant removals

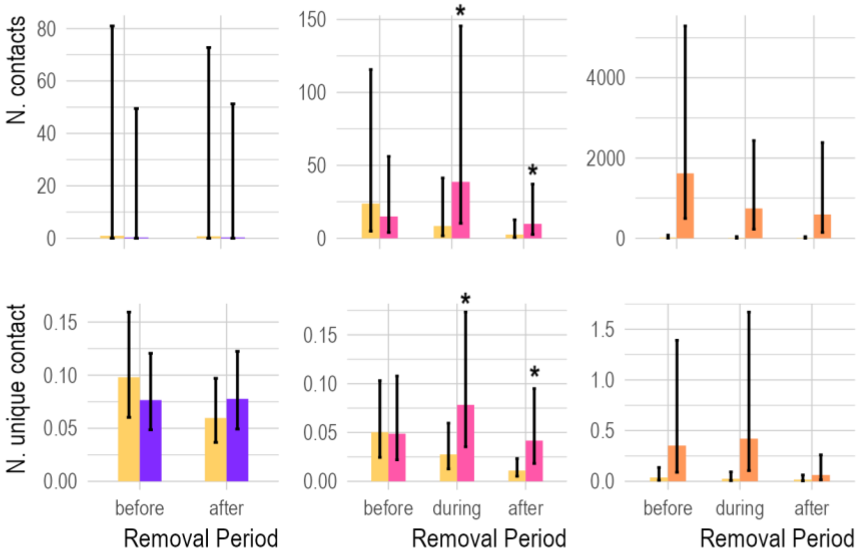

**Supplementary Figure 5.** Model predictions for contact responses by removal period (before- before removal activities occurred; during- during active removal activities; after- after removal activities ended), respective to each removal method, including the interaction of sex. *- interaction between removal treatment and removal period (i.e., effect of removal relative to control) was significant. **-interaction between removal treatment, period, and sex was significant. Error bars indicate 95% confidence interval. N. contacts - total number of contacts per day; N. unique contacts - number of unique individuals contacted per day; yellow-control; purple-aerial removals; pink-trapping removals; orange-toxicant removals

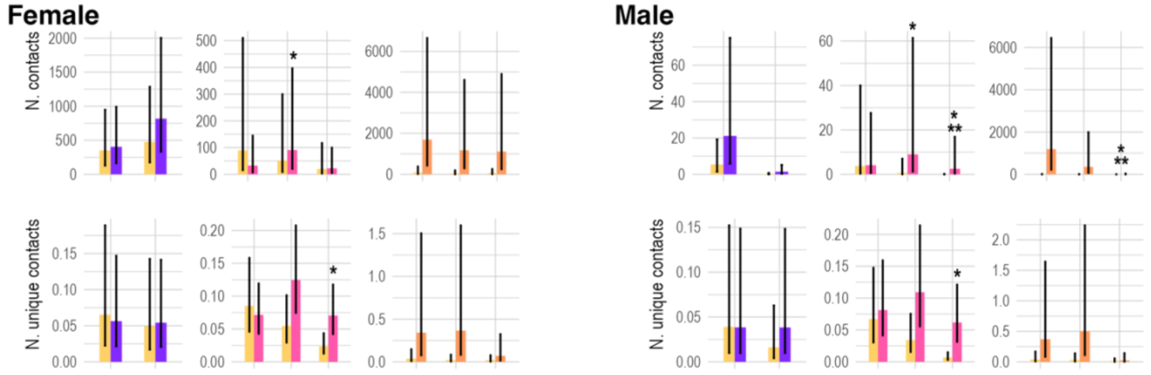

**Supplementary Figure 6.** Subsampling sensitivity analysis dotplot, demonstrating autocorrelated kernel density (akde) area estimates with 95% confidence intervals. A set of individuals with high resolution tracking data (15 minute fix rate) was subsampled to 240 minutes (4 hours) to determine whether area estimations from lower-resolution subsampled tracks diverged from estimates using the full track. Full- full, 15 minute resolution track; 240min- full track subsampled to 240 minute intervals.

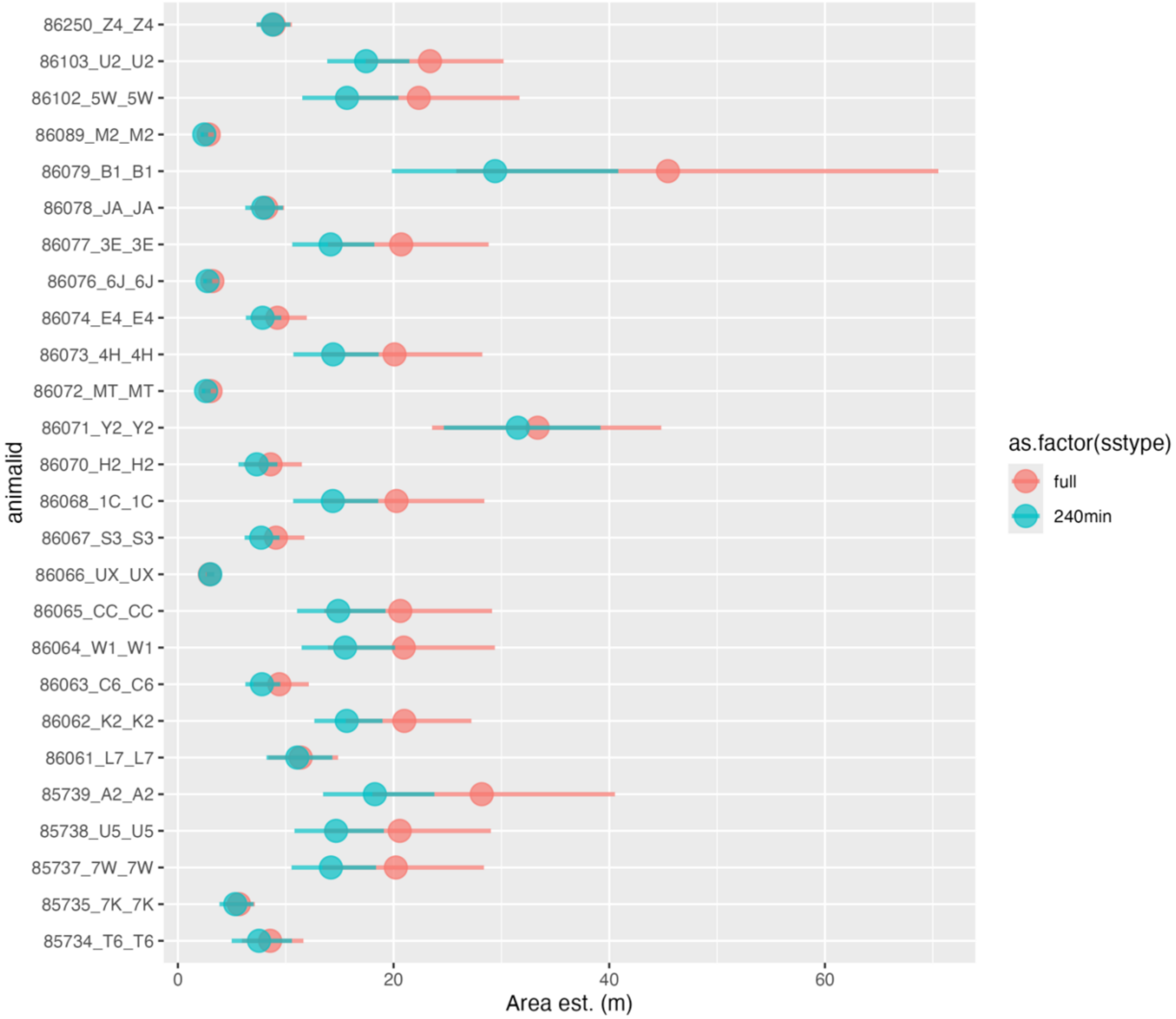

**Supplementary Figure 7.** Subsampling sensitivity analysis dotplot, demonstrating autocorrelated kernel density (akde) area estimates with 95% confidence intervals. A set of individuals with high resolution tracking data (15 minute fix rate) was subsampled to 20 minutes to determine whether area estimations from lower-resolution subsampled tracks diverged from estimates using the full track. Full- full, 15 minute resolution track; 20min- full track subsampled to 20 minute intervals.

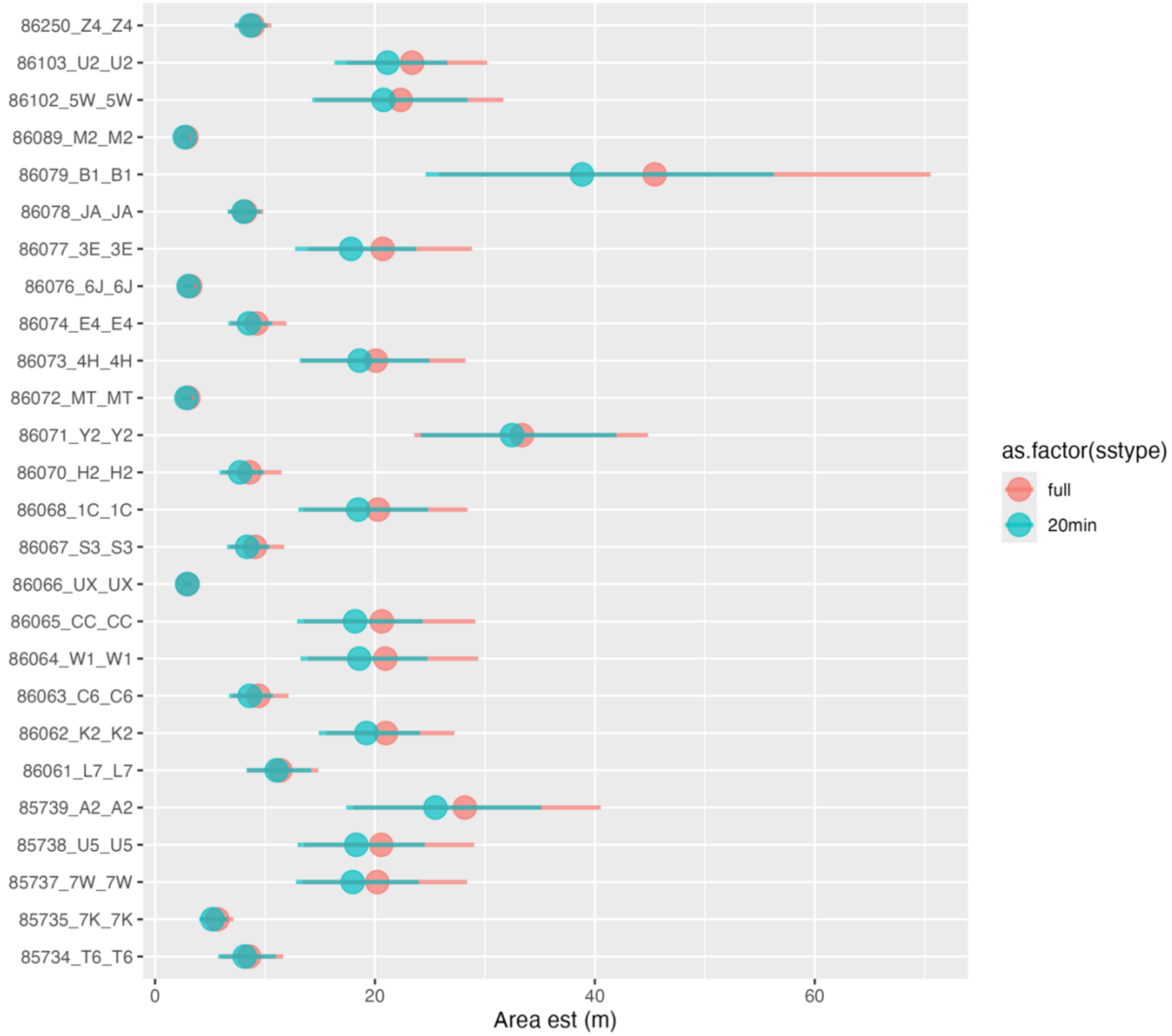

**Supplementary Figure 8.** Frequency distribution of differences in weekly displacement (m) between full and subsampled tracks (240 minute/4 hour intervals).

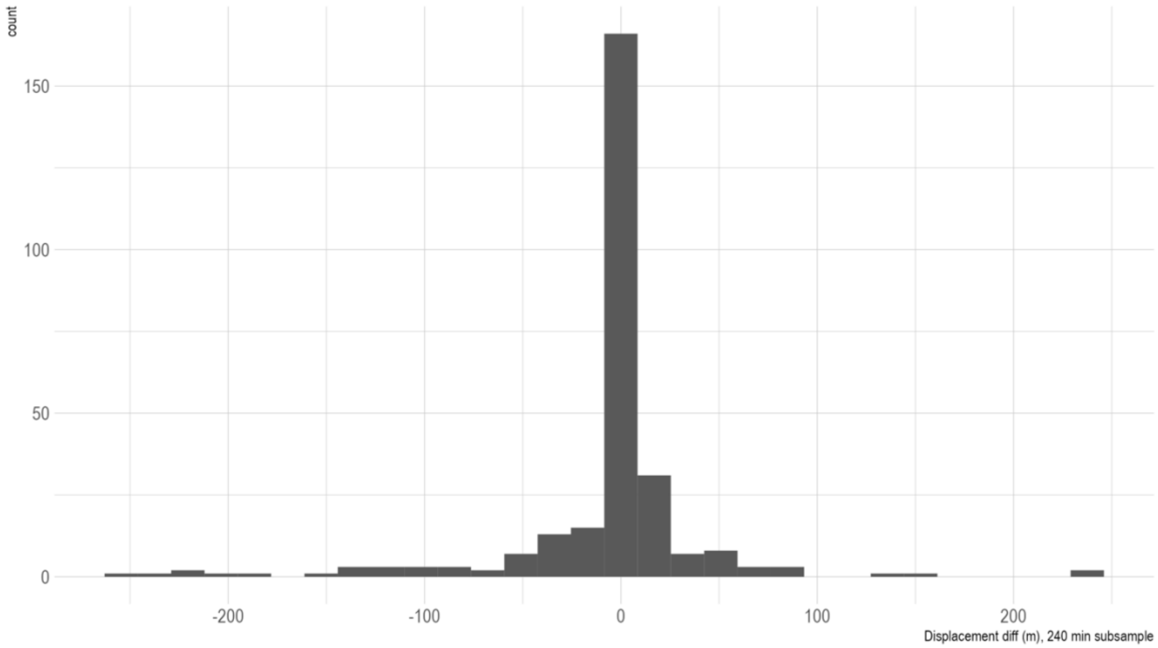

**Supplementary Figure 9.** Frequency distribution of differences in weekly displacement (m) between full and subsampled tracks (20 minute intervals).

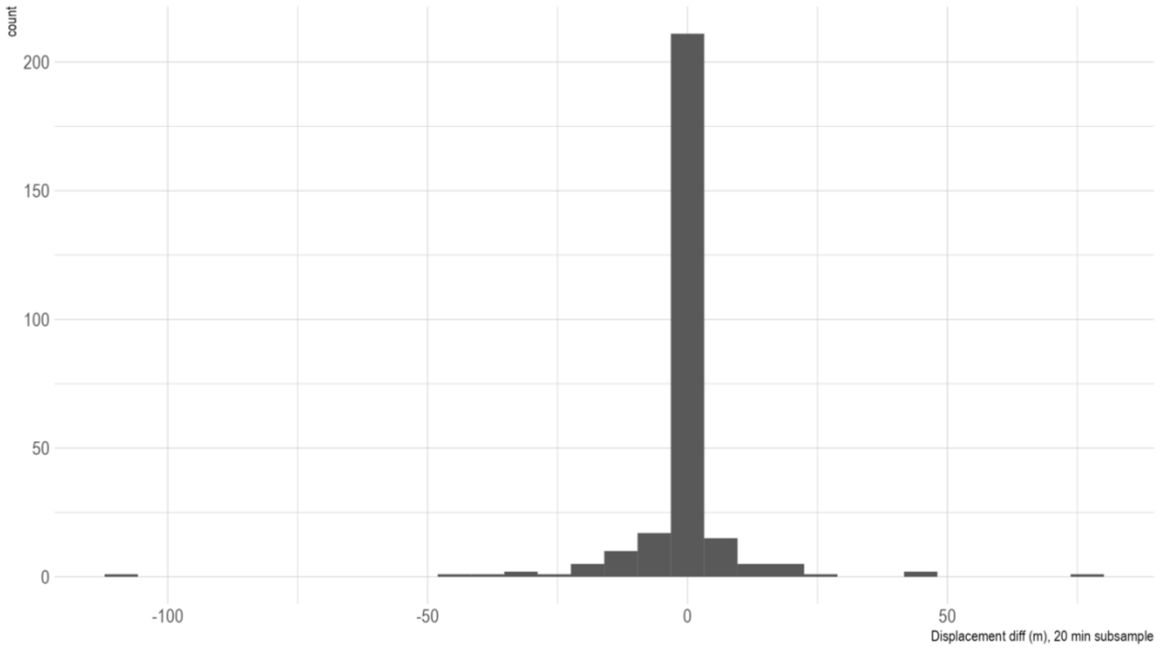

**Supplementary Figure 10.** Dot plot for sensitivity analysis on home range area responses of the toxicant treatment and control data, conducted for the significant interaction parameter between Removal type: tox and period: after. Overlap of a confidence interval with the dotted red x intercept indicates that the parameter is no longer significant when the respective individual is removed from the dataset. Individual removed- the individual identification number of the pig removed from each model.

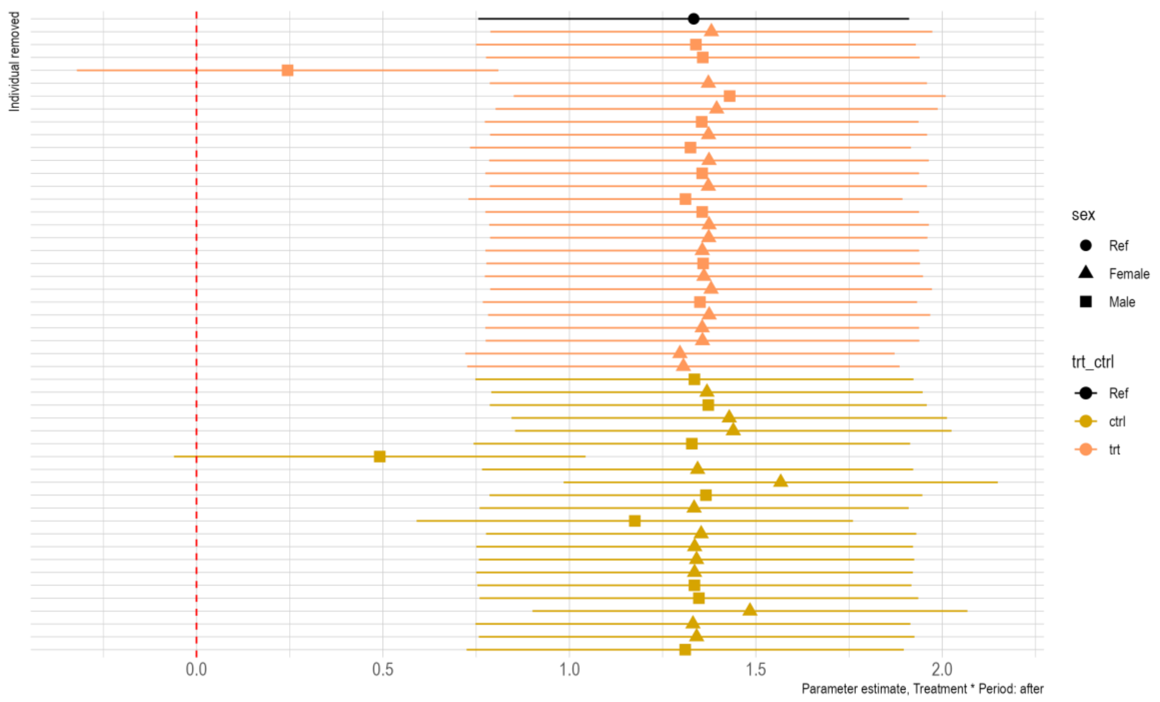

**Supplementary Figure 11.** Dot plot for sensitivity analysis on home range area responses of the toxicant treatment and control data, conducted for the significant interaction parameter between Removal type: tox and period: after. Overlap of a confidence interval with the dotted red x intercept indicates that the parameter is no longer significant when the respective individual is removed from the dataset. Individual removed- the individual identification number of the pig removed from each model.

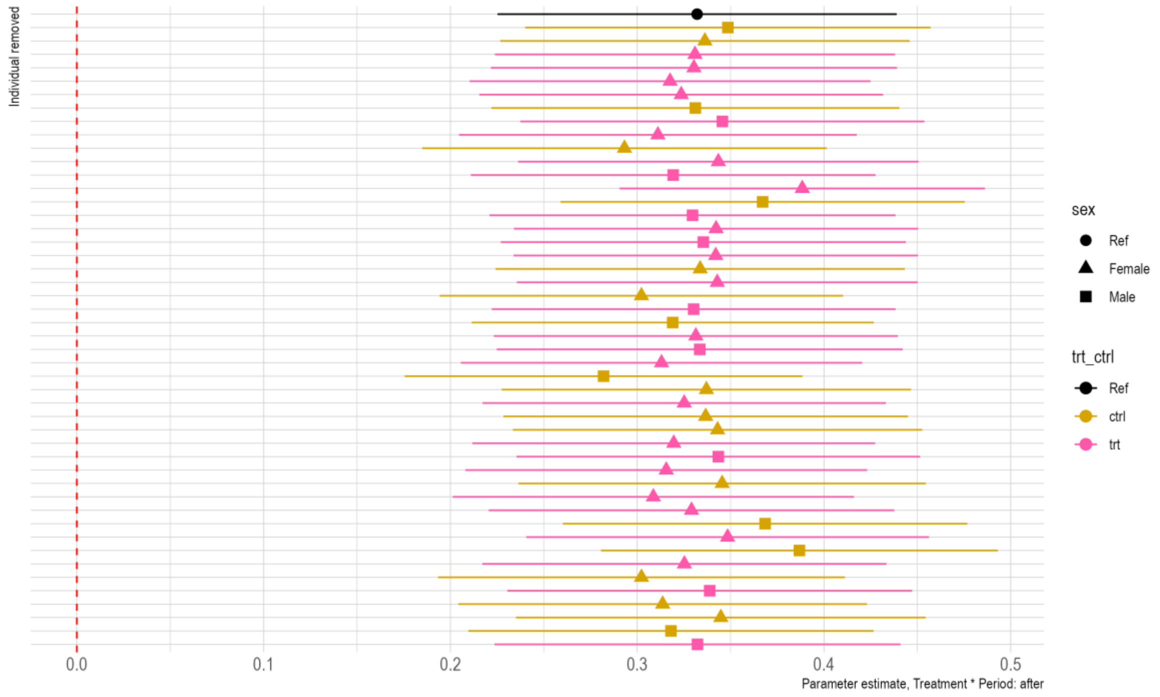

**Supplementary Figure 12.** Comparison of parameter estimates for each movement response modeled with the full dataset (full) and the dataset with pigs that died of toxicant bait removed (survivors). Periodafter- period following toxicant removals; periodduring- period during toxicant deployment; trt_ctrltrt- toxicant deployment treatment; trt_ctrltrt:periodafter- interaction between treatment and removal period; trt_ctrltrt:periodduring.

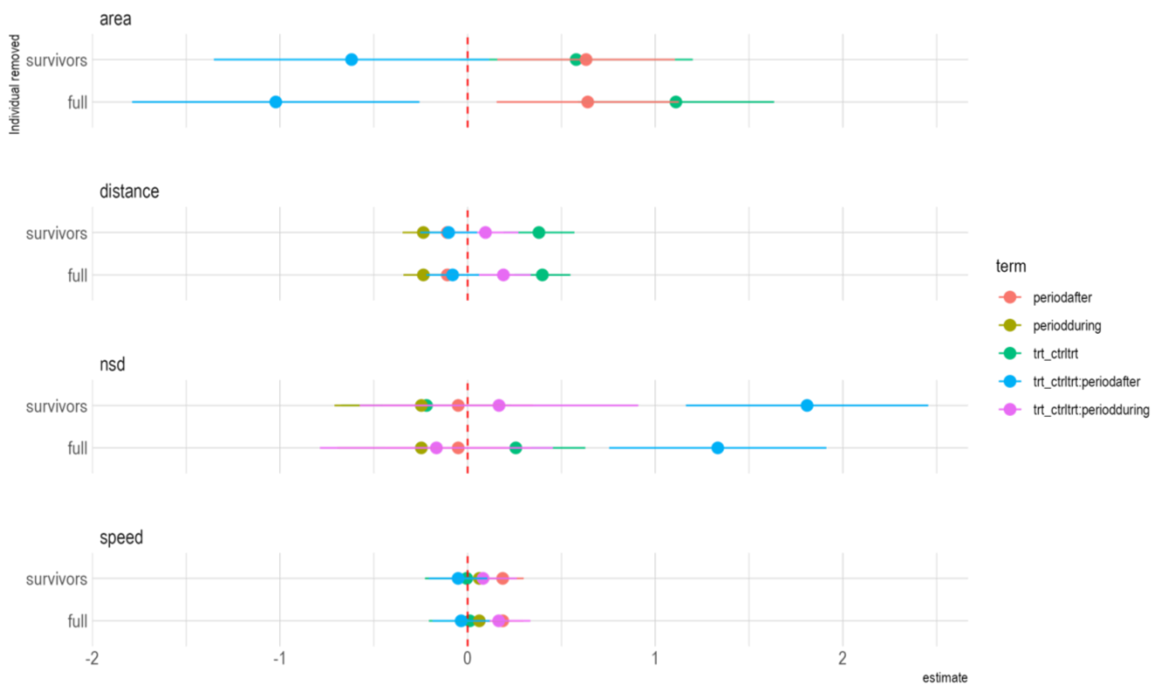

**Supplementary Table 1.** Parameter table for all models comparing movement response to removal treatment and period in a BACI design study on the effects of different removal operations on wild pigs. $p < 0.05$ was considered significant. Control individuals and the period before treatments began are the reference groups. Aerial- aerial treatments; tox- toxicant bait treatments; trap- trapping treatments. Exp(Beta)- exponentiated parameter estimate. Trt_ctrl- treatment or control; trt- treatment; ctrl- control.

| | aerial | | | tox | | | trap | | |
|---|---|---|---|---|---|---|---|---|---|
| **Characteristic** | **exp(Beta)** | **95% CI[1]** | **p-value** | **exp(Beta)** | **95% CI[1]** | **p-value** | **exp(Beta)** | **95% CI[1]** | **p-value** |
| **Area (km^2)** | | | | | | | | | |
| **trt_ctrl** | | | | | | | | | |
| ctrl | — | — | | — | — | | — | — | |
| trt | 0.74 | 0.36, 1.55 | 0.4 | 3.04 | 1.82, 5.07 | <0.001 | 0.74 | 0.34, 1.59 | 0.4 |
| **period** | | | | | | | | | |
| before | — | — | | — | — | | — | — | |
| after | 1.09 | 0.53, 2.24 | 0.8 | 1.90 | 1.18, 3.05 | 0.008 | 1.12 | 0.49, 2.59 | 0.8 |
| during | | | | | | | 1.09 | 0.47, 2.51 | 0.8 |
| **trt_ctrl * period** | | | | | | | | | |
| trt * after | 1.21 | 0.45, 3.29 | 0.7 | 0.36 | 0.17, 0.76 | 0.008 | 1.74 | 0.58, 5.17 | 0.3 |
| trt * during | | | | | | | 1.40 | 0.47, 4.16 | 0.6 |
| **NSD (m^2)** | | | | | | | | | |
| **trt_ctrl** | | | | | | | | | |
| ctrl | — | — | | — | — | | — | — | |
| trt | 0.84 | 0.36, 1.96 | 0.7 | 1.29 | 0.90, 1.86 | 0.2 | 1.08 | 0.44, 2.68 | 0.9 |
| **period** | | | | | | | | | |
| before | — | — | | — | — | | — | — | |
| after | 1.33 | 0.80, 2.21 | 0.3 | 0.95 | 0.65, 1.39 | 0.8 | 1.62 | 1.13, 2.33 | 0.008 |
| during | | | | 0.78 | 0.50, 1.21 | 0.3 | 1.46 | 1.10, 1.94 | 0.008 |
| **trt_ctrl * period** | | | | | | | | | |
| trt * after | 0.92 | 0.46, 1.84 | 0.8 | 3.79 | 2.15, 6.69 | <0.001 | 0.84 | 0.53, 1.33 | 0.5 |
| trt * during | | | | 0.85 | 0.46, 1.56 | 0.6 | 1.05 | 0.73, 1.50 | 0.8 |
| **distance (km)** | | | | | | | | | |
| **trt_ctrl** | | | | | | | | | |
| ctrl | — | — | | — | — | | — | — | |
| trt | 1.19 | 0.89,1.58 | 0.2 | 1.59 | 1.2,2.1 | <0.001 | 0.93 | 0.78, 1.12 | 0.5 |
| **period** | | | | | | | | | |
| before | — | — | | — | — | | — | — | |
| after | 1.13 | 1.03,1.23 | 0.008 | 0.87 | 0.8,0.95 | <0.001 | 0.99 | 0.91, 1.08 | 0.9 |
| during | | | | 0.81 | 0.74,0.89 | <0.001 | 0.90 | 0.83, 0.98 | 0.015 |
| **trt_ctrl * period** | | | | | | | | | |
| trt * after | 0.99 | 0.88,1.12 | 0.9 | 0.94 | 0.81,1.09 | 0.4 | 1.35 | 1.21, 1.50 | <0.001 |
| trt * during | | | | 1.28 | 1.12,1.47 | <0.001 | 1.68 | 1.51, 1.87 | <0.001 |
| **speed (km/hr)** | | | | | | | | | |
| **trt_ctrl** | | | | | | | | | |
| ctrl | — | — | | — | — | | — | — | |
| trt | 1.07 | 0.80, 1.44 | 0.7 | 1.03 | 0.7,1.51 | 0.9 | 0.95 | 0.73, 1.24 | 0.7 |
| **period** | | | | | | | | | |
| before | — | — | | — | — | | — | — | |
| after | 1.34 | 1.15, 1.57 | <0.001 | 0.97 | 0.77,1.21 | 0.8 | 1.28 | 1.05, 1.54 | 0.012 |
| during | | | | 0.94 | 0.76,1.17 | 0.6 | 1.18 | 0.98, 1.43 | 0.089 |
| **trt_ctrl * period** | | | | | | | | | |
| trt * after | 0.79 | 0.63, 0.98 | 0.032 | 1.21 | 0.84,1.75 | 0.3 | 0.94 | 0.73, 1.20 | 0.6 |
| trt * during | | | | 1.31 | 0.95,1.81 | 0.094 | 1.05 | 0.82, 1.35 | 0.7 |

[1]CI = Confidence Interval

**Supplementary Table 2.** Parameter table for all models comparing movement response to removal treatment and period in a BACI design study on the effects of different removal operations on wild pigs, with the interaction of sex. $p < 0.05$ was considered significant. Control females and the period before treatments began are the reference groups. Aerial- aerial treatments; tox- toxicant bait treatments; trap- trapping treatments. Exp(Beta)- exponentiated parameter estimate. Trt_ctrl- treatment or control; trt- treatment; ctrl- control.

| Characteristic | aerial | | | tox | | | trap | | |
|---|---|---|---|---|---|---|---|---|---|
| | exp(Beta) | 95% CI[1] | p-value | exp(Beta) | 95% CI[1] | p-value | exp(Beta) | 95% CI[1] | p-value |
| **NSD (m2)** | | | | | | | | | |
| **trt_ctrl** | | | | | | | | | |
| ctrl | — | — | | — | — | | — | — | |
| trt | 0.63 | 0.27, 1.52 | 0.3 | 3.28 | 1.31, 8.25 | 0.012 | 2.21 | 0.71, 6.85 | 0.2 |
| **period** | | | | | | | | | |
| before | — | — | | — | — | | — | — | |
| after | 2.38 | 1.72, 3.30 | <0.001 | 1.80 | 1.32, 2.46 | <0.001 | 2.84 | 1.70, 4.74 | <0.001 |
| during | | | | 1.44 | 1.05, 1.99 | 0.025 | 2.53 | 1.70, 3.79 | <0.001 |
| **sex** | | | | | | | | | |
| sexMale | 3.56 | 1.29, 9.81 | 0.014 | 2.75 | 0.98, 7.75 | 0.056 | 2.30 | 0.61, 8.76 | 0.2 |
| **trt_ctrl * period** | | | | | | | | | |
| trt * after | 0.80 | 0.51, 1.25 | 0.3 | 0.59 | 0.38, 0.91 | 0.018 | 0.34 | 0.17, 0.65 | 0.001 |
| trt * during | | | | 0.36 | 0.23, 0.57 | <0.001 | 0.44 | 0.26, 0.72 | 0.001 |
| **trt_ctrl * sex** | | | | | | | | | |
| trt * sexMale | 0.89 | 0.21, 3.74 | 0.9 | 1.12 | 0.28, 4.49 | 0.9 | 0.44 | 0.08, 2.63 | 0.4 |
| **period * sex** | | | | | | | | | |
| after * sexMale | 0.38 | 0.22, 0.64 | <0.001 | 0.73 | 0.45, 1.18 | 0.2 | 0.98 | 0.43, 2.22 | >0.9 |
| during * sexMale | | | | 0.71 | 0.44, 1.16 | 0.2 | 0.69 | 0.37, 1.30 | 0.3 |
| **trt_ctrl * period * sex** | | | | | | | | | |
| trt * after * sexMale | 1.44 | 0.68, 3.07 | 0.3 | 3.23 | 0.92, 11.3 | 0.066 | 3.71 | 1.25, 11.0 | 0.018 |
| trt * during * sexMale | | | | 2.65 | 1.31, 5.35 | 0.007 | 4.47 | 1.95, 10.2 | <0.001 |
| **Area (km2)** | | | | | | | | | |
| **trt_ctrl** | | | | | | | | | |
| ctrl | — | — | | — | — | | | | |
| trt | 0.75 | 0.31, 1.79 | 0.5 | 3.24 | 1.76, 5.98 | <0.001 | | | |
| **period** | | | | | | | | | |
| before | — | — | | — | — | | | | |
| after | 1.70 | 0.70, 4.10 | 0.2 | 2.63 | 1.49, 4.66 | <0.001 | | | |
| **sex** | | | | | | | | | |
| sexMale | 2.37 | 0.86, 6.55 | 0.10 | 2.49 | 1.23, 5.04 | 0.011 | | | |
| **trt_ctrl * period** | | | | | | | | | |
| trt * after | 0.63 | 0.19, 2.03 | 0.4 | 0.25 | 0.11, 0.55 | <0.001 | | | |
| **trt_ctrl * sex** | | | | | | | | | |
| trt * sexMale | 1.07 | 0.27, 4.23 | >0.9 | 0.87 | 0.34, 2.25 | 0.8 | | | |
| **period * sex** | | | | | | | | | |
| after * sexMale | 0.37 | 0.10, 1.44 | 0.2 | 0.51 | 0.21, 1.22 | 0.13 | | | |
| **trt_ctrl * period * sex** | | | | | | | | | |
| trt * after * sexMale | 4.19 | 0.61, 28.8 | 0.15 | 3.52 | 0.83, 14.9 | 0.088 | | | |
| **distance (km)** | | | | | | | | | |
| **trt_ctrl** | | | | | | | | | |
| ctrl | — | — | | — | — | | — | — | |
| trt | 1.13 | 0.96, 1.34 | 0.2 | 1.47 | 1.26, 1.71 | <0.001 | 0.87 | 0.73, 1.05 | 0.2 |
| **period** | | | | | | | | | |
| before | — | — | | — | — | | — | — | |
| after | 1.18 | 1.05, 1.33 | 0.005 | 0.85 | 0.76, 0.96 | 0.007 | 0.87 | 0.78, 0.97 | 0.015 |
| during | | | | 0.67 | 0.59, 0.77 | <0.001 | 0.89 | 0.80, 0.99 | 0.039 |
| **sex** | | | | | | | | | |
| Female | — | — | | — | — | | — | — | |
| Male | 1.51 | 1.25, 1.84 | <0.001 | 1.15 | 0.96, 1.37 | 0.13 | 1.25 | 1.00, 1.57 | 0.055 |

| Characteristic | aerial | | | tox | | | trap | | |
|---|---|---|---|---|---|---|---|---|---|
| | exp(Beta) | 95% CI[1] | p-value | exp(Beta) | 95% CI[1] | p-value | exp(Beta) | 95% CI[1] | p-value |
| **trt_ctrl * period** | | | | | | | | | |
| trt * after | 0.82 | 0.70, 0.96 | 0.011 | 0.88 | 0.75, 1.04 | 0.13 | 1.57 | 1.36, 1.80 | <0.001 |
| trt * during | | | | 1.25 | 1.05, 1.50 | 0.014 | 1.75 | 1.52, 2.01 | <0.001 |
| **trt_ctrl * sex** | | | | | | | | | |
| trt * Male | 1.37 | 1.04, 1.79 | 0.025 | 1.04 | 0.82, 1.31 | 0.8 | 1.31 | 0.97, 1.77 | 0.080 |
| **period * sex** | | | | | | | | | |
| after * Male | 0.87 | 0.72, 1.04 | 0.12 | 1.13 | 0.95, 1.35 | 0.2 | 1.34 | 1.14, 1.59 | <0.001 |
| during * Male | | | | 1.41 | 1.15, 1.74 | <0.001 | 1.03 | 0.87, 1.21 | 0.8 |
| **trt_ctrl * period * sex** | | | | | | | | | |
| trt * after * Male | 1.06 | 0.83, 1.37 | 0.6 | 1.37 | 1.02, 1.84 | 0.039 | 0.70 | 0.56, 0.88 | 0.002 |
| trt * during * Male | | | | 0.98 | 0.74, 1.30 | 0.9 | 0.90 | 0.72, 1.12 | 0.4 |
| **speed (km/hr)** | | | | | | | | | |
| **trt_ctrl** | | | | | | | | | |
| ctrl | — | — | | — | — | | — | — | |
| trt | 0.93 | 0.70, 1.22 | 0.6 | 1.47 | 1.26, 1.71 | <0.001 | 0.77 | 0.57, 1.03 | 0.076 |
| **period** | | | | | | | | | |
| before | — | — | | — | — | | — | — | |
| after | 1.45 | 1.18, 1.78 | <0.001 | 0.85 | 0.76, 0.96 | 0.007 | 1.19 | 0.93, 1.51 | 0.2 |
| during | | | | 0.67 | 0.59, 0.77 | <0.001 | 1.01 | 0.79, 1.28 | >0.9 |
| **sex** | | | | | | | | | |
| Female | — | — | | — | — | | — | — | |
| Male | 1.41 | 1.03, 1.93 | 0.034 | 1.15 | 0.96, 1.37 | 0.13 | 0.84 | 0.59, 1.20 | 0.3 |
| **trt_ctrl * period** | | | | | | | | | |
| trt * after | 0.69 | 0.52, 0.90 | 0.007 | 0.88 | 0.75, 1.04 | 0.13 | 0.97 | 0.72, 1.31 | 0.8 |
| trt * during | | | | 1.25 | 1.05, 1.50 | 0.014 | 1.09 | 0.81, 1.48 | 0.6 |
| **trt_ctrl * sex** | | | | | | | | | |
| trt * Male | 1.73 | 1.11, 2.69 | 0.016 | 1.04 | 0.82, 1.31 | 0.8 | 1.86 | 1.15, 2.98 | 0.011 |
| **period * sex** | | | | | | | | | |
| after * Male | 0.85 | 0.62, 1.16 | 0.3 | 1.13 | 0.95, 1.35 | 0.2 | 1.19 | 0.82, 1.72 | 0.4 |
| during * Male | | | | 1.41 | 1.15, 1.74 | <0.001 | 1.44 | 1.00, 2.08 | 0.053 |
| **trt_ctrl * period * sex** | | | | | | | | | |
| trt * after * Male | 1.37 | 0.89, 2.13 | 0.2 | 1.37 | 1.02, 1.84 | 0.039 | 0.95 | 0.58, 1.55 | 0.8 |
| trt * during * Male | | | | 0.98 | 0.74, 1.30 | 0.9 | 0.97 | 0.59, 1.58 | 0.9 |

[1]CI = Confidence Interval

**Supplementary Table 3.** Parameter table for all models comparing contact responses to removal treatment and period in a BACI design study on the effects of different removal operations on wild pigs. $p < 0.05$ was considered significant. Control individuals and the period before treatments began are the reference groups. Aerial- aerial treatments; tox- toxicant bait treatments; trap- trapping treatments. Exp(Beta)- exponentiated parameter estimate. Trt_ctrl- treatment or control; trt- treatment; ctrl- control.

| | tox | | | trap | | | aer | | |
|---|---|---|---|---|---|---|---|---|---|
| **Characteristic** | **exp(Beta)** | **95% CI[1]** | **p-value** | **exp(Beta)** | **95% CI[1]** | **p-value** | **exp(Beta)** | **95% CI[1]** | **p-value** |
| **N total contacts** | | | | | | | | | |
| **trt_ctrl** | | | | | | | | | |
| ctrl | — | — | | — | — | | — | — | |
| trt | 39.5 | 3.37, 463 | 0.003 | 1.19 | 0.06,25.52 | >0.9 | 0.33 | 0,229.03 | 0.7 |
| **period** | | | | | | | | | |
| before | — | — | | — | — | | — | — | |
| during | 0.50 | 0.16, 1.56 | 0.2 | 0.42 | 0.15,1.18 | 0.094 | | | |
| after | 0.49 | 0.17, 1.37 | 0.2 | 0.13 | 0.04,0.39 | <0.001 | 0.80 | 0.12,5.53 | 0.8 |
| **trt_ctrl * period** | | | | | | | | | |
| trt * during | 0.54 | 0.09, 3.11 | 0.5 | 5.91 | 1.52,22.94 | 0.009 | | | |
| trt * after | 1.02 | 0.18, 5.85 | >0.9 | 4.88 | 1.23,19.35 | 0.021 | 1.31 | 0.13,13.45 | 0.8 |
| **N unique contacts** | | | | | | | | | |
| **trt_ctrl** | | | | | | | | | |
| ctrl | — | — | | — | — | | — | — | |
| trt | 5.25 | 0.97,28.23 | 0.049 | 0.97 | 0.36,2.65 | >0.9 | 0.78 | 0.40, 1.52 | 0.5 |
| **period** | | | | | | | | | |
| before | — | — | | — | — | | — | — | |
| during | 0.68 | 0.34,1.38 | 0.3 | 0.55 | 0.28,1.08 | 0.077 | | | |
| after | 0.46 | 0.22,0.95 | 0.032 | 0.22 | 0.11,0.44 | <0.001 | 0.61 | 0.31, 1.21 | 0.2 |
| **trt_ctrl * period** | | | | | | | | | |
| trt * during | 1.48 | 0.5,4.39 | 0.5 | 2.93 | 1.21,7.12 | 0.015 | | | |
| trt * after | 0.56 | 0.18,1.72 | 0.3 | 3.90 | 1.61,9.48 | 0.002 | 1.67 | 0.65, 4.28 | 0.3 |

[1]CI = Confidence Interval

**Supplementary Table 4.** Parameter table for all models comparing contact responses to removal treatment and period in a BACI design study on the effects of different removal operations on wild pigs, with the interaction of sex. $p < 0.05$ was considered significant. Control females and the period before treatments began are the reference groups. Aerial- aerial treatments; tox- toxicant bait treatments; trap- trapping treatments. Exp(Beta)- exponentiated parameter estimate. Trt_ctrl- treatment or control; trt- treatment; ctrl- control.

| | tox | | | trap | | | aer | | |
|---|---|---|---|---|---|---|---|---|---|
| **Characteristic** | **exp(Beta)** | **95% CI[1]** | **p-value** | **exp(Beta)** | **95% CI[1]** | **p-value** | **exp(Beta)** | **95% CI[1]** | **p-value** |
| **N total contacts** | | | | | | | | | |
| **trt_ctrl** | | | | | | | | | |
| ctrl | — | — | | — | — | | | | |
| trt | 83.0 | 0.71,9679.18 | 0.063 | 0.37 | 0.04, 3.55 | 0.4 | | | |
| **period** | | | | | | | | | |
| before | — | — | | — | — | | | | |
| during | 0.42 | 0.11,1.68 | 0.2 | 0.57 | 0.19, 1.74 | 0.3 | | | |
| after | 0.60 | 0.18,2.05 | 0.4 | 0.22 | 0.07, 0.71 | 0.011 | | | |
| **sex** | | | | | | | | | |
| sexMale | 0.05 | 0,0.85 | 0.034 | 0.04 | 0.00, 0.78 | 0.034 | | | |
| **trt_ctrl * period** | | | | | | | | | |
| trt * during | 1.31 | 0.19,8.83 | 0.8 | 4.81 | 1.08, 21.4 | 0.039 | | | |
| trt * after | 2.05 | 0.34,12.5 | 0.4 | 3.09 | 0.67, 14.1 | 0.15 | | | |
| **trt_ctrl * sex** | | | | | | | | | |
| trt * sexMale | 36.2 | 0.13,10129.32 | 0.2 | 2.95 | 0.07, 129 | 0.6 | | | |
| **period * sex** | | | | | | | | | |
| during * sexMale | 1.72 | 0.16,18.43 | 0.6 | 0.28 | 0.03, 2.84 | 0.3 | | | |
| after * sexMale | 0.28 | 0.03,2.71 | 0.3 | 0.02 | 0.00, 0.22 | 0.002 | | | |
| **trt_ctrl * period * sex** | | | | | | | | | |
| trt * during * sexMale | 0.05 | 0,2.55 | 0.13 | 2.81 | 0.16, 48.5 | 0.5 | | | |
| trt * after * sexMale | 0.00 | 0,0.28 | 0.012 | 54.3 | 2.55, 1,153 | 0.010 | | | |
| **N unique contacts** | | | | | | | | | |
| **trt_ctrl** | | | | | | | | | |
| ctrl | — | — | | — | — | | — | — | |
| trt | 4.46 | 0.49,40.7 | 0.2 | 0.84 | 0.38, 1.87 | 0.7 | 0.00 | 0,0 | <0.001 |
| **period** | | | | | | | | | |
| before | — | — | | — | — | | — | — | |
| during | 0.54 | 0.23,1.27 | 0.2 | 0.64 | 0.27, 1.54 | 0.3 | | | |
| after | 0.52 | 0.22,1.23 | 0.13 | 0.28 | 0.11, 0.66 | 0.004 | 1.28 | 0.65,2.53 | 0.5 |
| **sex** | | | | | | | | | |
| sexMale | 1.03 | 0.27,3.97 | >0.9 | 0.79 | 0.29, 2.14 | 0.6 | 0.64 | 0.23,1.76 | 0.4 |
| **trt_ctrl * period** | | | | | | | | | |
| trt * during | 1.74 | 0.5,6.05 | 0.4 | 2.72 | 0.87, 8.45 | 0.085 | | | |
| trt * after | 0.62 | 0.18,2.13 | 0.4 | 3.59 | 1.15, 11.2 | 0.027 | 0.69 | 0.27,1.72 | 0.4 |
| **trt_ctrl * sex** | | | | | | | | | |
| trt * sexMale | 2.72 | 0.17,42.13 | 0.5 | 1.44 | 0.39, 5.36 | 0.6 | 3.65 | 0.89,14.89 | 0.066 |
| **period * sex** | | | | | | | | | |
| during * sexMale | 1.52 | 0.39,5.95 | 0.5 | 0.80 | 0.19, 3.28 | 0.8 | | | |
| after * sexMale | 0.61 | 0.15,2.52 | 0.5 | 0.37 | 0.09, 1.52 | 0.2 | 0.52 | 0.17,1.63 | 0.3 |
| **trt_ctrl * period * sex** | | | | | | | | | |
| trt * during * sexMale | 0.69 | 0.06,7.47 | 0.8 | 0.97 | 0.15, 6.19 | >0.9 | | | |
| trt * after * sexMale | 0.18 | 0.01,3.91 | 0.3 | 2.09 | 0.33, 13.3 | 0.4 | 2.50 | 0.56,11.22 | 0.2 |

[1]CI = Confidence Interval
